# Regulation of de- and reciliation by KRAS during muscle cell differentiation

**DOI:** 10.64898/2026.07.07.736926

**Authors:** Rohan Chippalkatti, Bianca Parisi, Elisabeth Schaffner-Reckinger, Christina Laurini, Atanasio Gómez-Mulas, Zoe Geimer, Daniel Kwaku Abankwa

## Abstract

The primary cilium has been implicated in multiple developmental processes, such as cell migration and asymmetric cell division of stem- and progenitor cells. While most in vitro model systems examine ciliogenesis induced by serum starvation, it is not fully understood how de-and re-ciliation are regulated in proliferating stem- and progenitor cells.

Here we employ the hierarchically organized C2C12 skeletal muscle cell line to examine how K-Ras4B participates in de- and re-ciliation processes of ciliated stem- and progenitor cells. We show that MAPK-pathway activation supports ciliogenesis through phosphorylation of centrosomal protein CEP55, which can then no longer stabilize the master regulator of de-ciliation Aurora kinase A. K-Ras4B localizes to the primary cilium aided by the ciliary trafficking chaperone PDE6D, which promotes ciliation. In line with this, depletion of components of the PDE6D machinery, RPGR and RPGRIP1L, decreases ciliation. Activation of the ciliary AMPK-PKG2-pathway increases S181-phosphorylation of K-Ras4B, which negatively regulates its binding to PDE6D, its ciliary abundance and promotes differentiation. Our work integrates a major mediator of mitogenic signaling into the regulation of ciliogenesis of proliferating muscle stem- and progenitor cells.

## Introduction

The primary cilium is a sensory organelle that is organized as a few micrometers long membrane protrusion with a microtubule bundle, the axoneme, at its core. This bundle emerges from the basal body, which becomes the mother centrosome during interphase. Primary cilia are found in most stem and progenitor cells, while also other cell types can form cilia (Yanardag & Pugacheva, 2021). By contrast, motile cilia possess a modified axoneme and are found on several terminally differentiated epithelia, such as of the kidney and uterus (Gopalakrishnan *et al*, 2023).

The primary cilium has multiple, interleaved sensory functions during development and tissue repair, such as coordination of cell migration and axonal guidance (Atkins *et al*, 2025; Stoufflet & Caille, 2022). Mechanical stimulation of the primary cilium can be registered via activation of ciliary LKB1, which phosphorylates Thr172 of AMPK on the basal body (Boehlke *et al*, 2010). This facilitates AMPK activation, which suppresses mTORC1 by AMPK-mediated phosphorylation and activation of Tsc2, a negative regulator of the pathway (Lai & Jiang, 2020). Several lines of evidence furthermore support the activity of the Ras-MAPK pathway in the cilium. Two FGF receptors, FGFR1 and FGFR2, localize to the primary cilium and activate the pathway to control FGF-target gene expression and ciliary length via the ICK kinase (Kunova Bosakova *et al*, 2019; Nita *et al*, 2025). In growth arrested fibroblasts, PDGFR⍺ is found in the cilium and, upon activation, MEK1/2 is phosphorylated within the cilium and on the basal body (Schneider *et al*, 2005). The most characteristic sensory functions of the cilium are executed by receptors of crucial developmental and stemness pathways such as Hedgehog, Wnt and Notch, which are usually confined to the primary cilium (Liu *et al*, 2018).

Another fundamental function that is associated with this developmental receptor repertoire is the ability of the primary cilium to confer stemness during tissue development (Yanardag & Pugacheva, 2021). When stem cells divide asymmetrically they give rise to one stem cell of equal potency (i.e. self-renewal), while the other daughter cell is committed to differentiate (Chen & Yamashita, 2021). Examples from neural and muscle stem cells suggest that during this process, the reformation of the primary cilium segregates with the cell that retains stemness properties (Chen & Yamashita, 2021; Jaafar Marican *et al*, 2016; Paridaen *et al*, 2013). This implies that de- and re-ciliation cycles during asymmetric cell division are critical for cell differentiation and development.

However, our current knowledge about this process is mostly derived from model systems, such as serum-starved NIH3T3 cells, where re-ciliation is induced as cells exit the cell cycle in G1 when serum is withdrawn, and not from proliferating stem- and progenitor cell systems (Gopalakrishnan *et al*., 2023). Critical ciliogenesis steps involve the removal of the centriolar coiled-coil protein 110 (CP110) that caps the mother centriole, which involves Tau tubulin kinase 2 (TTBK2). CEP164 engages Rab8 and ciliary vesicle formation, while residual activity of the master regulator of deciliation, Aurora kinase A, is suppressed mostly by inactivating its activators (Korobeynikov *et al*, 2017).

Conversely, cilium disassembly at G1/S transition is initiated by active Aurora kinase A, which integrates multiple signals, notably the growth factor stimulated increase of scaffold protein NEDD9 (Korobeynikov *et al*., 2017; Pugacheva *et al*, 2007). NEDD9 facilitates Aurora kinase A activation and together with PLK1 triggers histone deacetylase 6 (HDAC6), which deacetylates ciliary microtubules, leading to their disassembly. Several additional molecular steps ultimately lead to the recruitment of centrosomal protein 97 (CEP97) and CP110 to the mother centriole, which caps it for cilium assembly as cells progress through the cell cycle (Korobeynikov *et al*., 2017).

Dysregulation of the primary cilium impacts not only on stemness and developmental signaling pathways but has the potential to perturb cell differentiation and thus tissue homeostasis and repair in the adult (Fu *et al*, 2014; Venugopal *et al*, 2020). In line with this, cilia are lost in most cancer cells leading to profoundly perturbed tissue differentiation in the form of a tumor (Liu *et al*., 2018). On the other hand, germline mutations of ciliary genes are associated with severe but rare developmental diseases collectively referred to as ciliopathies (Reiter & Leroux, 2017). Ciliopathies can affect individual ciliated organs, such as the kidney, or multiple organ systems (brain, heart, muscle etc.).

Many ciliopathy genes regulate ciliary access of proteins, as diffusive entry into the cilium is only possible for proteins <70 kDa (Endicott & Brueckner, 2018). To this end, lipid-modified proteins require trafficking chaperones such as PDE6D, which release cargo via allosteric release factors Arl2 and Arl3 (Fansa & Wittinghofer, 2016; Ismail *et al*, 2011). When in the active, GTP-bound state, Arl2 can bind to PDE6D and facilitate the release of cargo such as Ras-proteins, which bind with only micromolar or submicromolar affinities to PDE6D (Fansa & Wittinghofer, 2016; Ismail *et al*., 2011). Engagement of PDE6D is determined by residues and post-translational modifications, such as palmitoylation, which sterically interfere with the entrance of the prenyl-binding pocket. Thus, neither K-Ras4A nor palmitoylated N-Ras and H-Ras are cargos of PDE6D, while K-Ras4B (**hereafter K-Ras**) is a micromolar binder (Chandra *et al*, 2011; Dharmaiah *et al*, 2016; Siddiqui *et al*, 2020). GTP-Arl2 mediated release of PDE6D-bound K-Ras4B in the perinuclear area permits K-Ras trapping on the recycling endosomes for vesicular forward trafficking to the plasma membrane (Schmick *et al*, 2014). However, plasma membrane delivery of K-Ras only partially depends on PDE6D (Kaya *et al*, 2024; Pavic *et al*, 2022), which may hint at a more specific function of PDE6D in conjunction with K-Ras.

The ciliary release machinery of PDE6D comprises the ciliary marker protein Arl13B, which is a guanine nucleotide exchange factor (GEF) of Arl3 (Gotthardt *et al*, 2015). In line with the fact that low affinity PDE6D cargo such as the Ras-related small GTPase Rheb is released by GTP-Arl3 from PDE6D (Fansa *et al*, 2016), we recently showed that K-Ras, but not N-Ras and H-Ras, localizes PDE6D-dependently to the membrane and basal body of the primary cilium of C2C12 skeletal muscle cells (Chippalkatti *et al*, 2026). The presence of B-Raf and active MEK on the basal body and of active ERK inside the cilium suggests that K-Ras is active there. Supported by mathematical modelling and single cell RNA transcriptomics, we inferred that ciliary K-Ras restricts commitment and thus protects stemness of muscle stem cells during their asymmetric cell divisions. High *KRAS* expression was associated with proliferating stem and progenitor cells, which undergo de- and re-ciliation cycles during asymmetric divisions, but the molecular pathways connecting ciliary K-Ras to these processes remained unresolved.

K-Ras regulates de- and re-ciliation of C2C12 skeletal muscle cells, which recapitulate essential steps of in vivo muscle differentiation (Chippalkatti *et al*, 2024; Chippalkatti *et al*., 2026). This cell line is hierarchically organized, with ∼1% muscle stem cells (**MuSC**; ∼77% ciliated) at the apex, which sustains via ciliation-guided asymmetric cell divisions the much larger population of transit amplifying cells (**TAC**; ∼13% ciliated) in high serum. Switching to low serum then induces differentiation, which progresses through intermediate TAC states toward terminal differentiation (Chippalkatti *et al*., 2026).

Here we describe how re-ciliation is supported by K-Ras-MAPK signaling, which increases S423/S426-phosphorylation of the centriolar protein CEP55 thus blocking its ability to engage de-ciliation stimulating Aurora kinase A. Conversely, de-ciliation is regulated via phosphorylation of Ser181 at the C-terminus of K-Ras downstream of the ciliary AMPK-PKG2-pathway, which reduces K-Ras association with PDE6D and thus its ciliary localization.

## Results

### MAPK-activity increases CEP55-phosphorylation which reduces its binding to AURKA

We recently described that K-Ras sustains ciliation during asymmetric cell divisions of C2C12 skeletal muscle cells and thus impacts on cell differentiation. We found that both K-Ras and MAPK-components localize to the basal body and inside the primary cilium, suggesting active MAPK-signaling there (Chippalkatti *et al*., 2026).

We therefore sought to identify the molecular target downstream of ciliary K-Ras-MAPK-signaling that could sustain ciliation. We assumed that a target protein would be localized to the basal body and associated with cancer and/ or developmental diseases. Our search converged on centrosomal protein 55 kDa (CEP55), which is associated with ciliopathies Joubert Syndrome 3 and Meckel-Gruber syndrome (MKS) (Bondeson *et al*, 2017; Frosk *et al*, 2017; Rawlins *et al*, 2019). CEP55 is upregulated in several tumors, regulates cytokinesis, genomic stability and stemness, and was reported to localize preferentially to the mother centriole (Fabbro *et al*, 2005; Jeffery *et al*, 2016; Kalimutho *et al*, 2018). We confirmed the centriolar localization in C2C12 cells, where EGFP-CEP55 localized to both centrioles with an apparent enrichment on the mother centriole/ basal body that gives rise to the cilium (**Figure 1A**).

**Figure 1.**
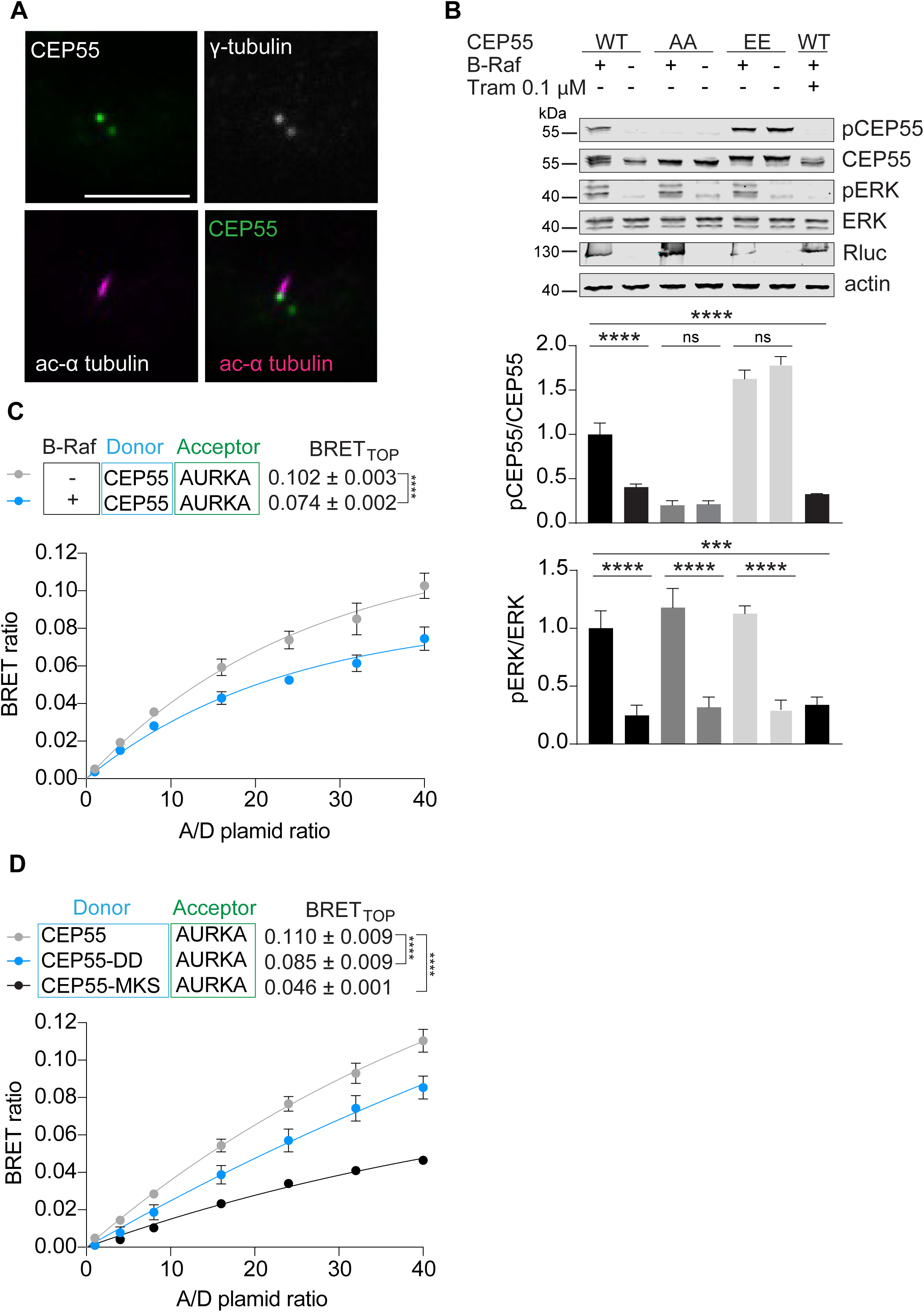
Phosphorylation of CEP55 downstream of the Ras-MAPK pathway disrupts its interaction with AURKA. (A) Confocal images of C2C12 cells transfected with EGFP-CEP55 and grown in high serum for 48 h. Cells were immunolabelled for centriolar/ basal body marker γ-tubulin (top) or acetylated α-tubulin marking the cilium (bottom). Scale bar = 5 µm. (B) Immunoblot analysis showing phosphorylation of CEP55 or ERK1/2 downstream of the MAPK pathway in HEK293 cells transfected with Rluc8-tagged B-Raf and Flag-tagged CEP55 or phospho-site S425/S428 mutants CEP55-AA or CEP55-EE. Shown are representative images of immunoblots labelled with phospho-CEP55, CEP55, phospho-ERK, ERK, Renilla luciferase or actin (loading control) antibodies. The plots show the quantification of relative CEP55 or ERK phosphorylation, N = 4, except for trametinib (Tram) treatment where N = 3. Means ± SEM are plotted. Statistical analysis was done with one-way ANOVA and Tukey’s multiple comparisons test. Shown are the statistical comparisons with matched control condition. (C) BRET titration curves of Rluc8-CEP55 and GFP2-AURKA expressed in HEK293 cells, with or without co-expression of mCherry-B-Raf (200 ng DNA). N = 3. Statistical analysis was performed with the extra sum-of-squares F-test. (D) BRET titration curves of Rluc8-CEP55 derived phospho-site mutant S423D/S426D (-DD), Rluc8-CEP55-MKS and GFP2-AURKA expressed in HEK293 cells, N = 3. Statistical analysis was performed with the extra sum-of-squares F-test.

CEP55 possesses one N-terminal and one C-terminal coiled-coil domain that are joined by the ESCRT and ALIX-binding region (Jeffery *et al*., 2016). The C-terminal fragment beyond the last coiled-coil domain, which corresponds to amino acids 355-462 of human CEP55, is sufficient to interact with and stabilize the master regulator of de-ciliation Aurora kinase A (**AURKA**) (Nishimura *et al*, 2021; Zhang *et al*, 2021). Two major phosphorylation sites at the C-terminus, S423/S426 (S425/S428 of the human protein), promote the translocation of CEP55 from the centrosome to the midbody and are the targets of ERK2 (Fabbro *et al*., 2005; Kalimutho *et al*., 2018).

We therefore hypothesized that ciliary K-Ras-MAPK-activity on the basal body would phosphorylate S423/S426 (corresponding to human S425/S428), thus disrupting the interaction of phospho-CEP55 with AURKA, which would become destabilized and could no longer promote disassembly of the cilium.

In line with this, co-expression of B-Raf significantly increased the phosphorylation of CEP55, while it had no effect on the human CEP55-S425E/S428E (CEP55-EE) mutant with phosphomimetic substitutions or non-phosphorylatable CEP55-S425A/S428A (CEP55-AA) mutant (**Figure 1B**). Likewise, co-expression of B-Raf significantly reduced the BRET-interaction of CEP55 with AURKA (**Figure 1C**), suggesting that phospho-CEP55 binds less to AURKA. This was further supported by BRET-interaction data showing that phosphomimetic CEP55-DD has a significantly reduced ability to bind to AURKA, albeit not as much as the C-terminal truncation mutant associated with MKS (CEP55-MKS, residues 1-85) (Bondeson *et al*., 2017) (**Figure 1D**).

Therefore, increased K-Ras-MAPK pathway activity could increase CEP55 phosphorylation and thus decrease its interaction with and stabilization of AURKA (Zhang *et al*., 2021).

### Phospho-CEP55 stabilizes ciliation of proliferating C2C12 cells

Consistent with these results, we found that siRNA-mediated knockdown of both K-Ras isoforms significantly increased AURKA levels in C2C12 cells (**Figure 2A**). In line with this effect being mediated by CEP55, co-expression of wt CEP55 with B-Raf decreased AURKA levels significantly and in dependence of MEK1/2 activity, as MEK1/2 inhibitor trametinib curtailed this response. Accordingly, AURKA levels were low with the expression of CEP55-EE, with or without co-expression of B-Raf (**Figure 2B**).

**Figure 2:**
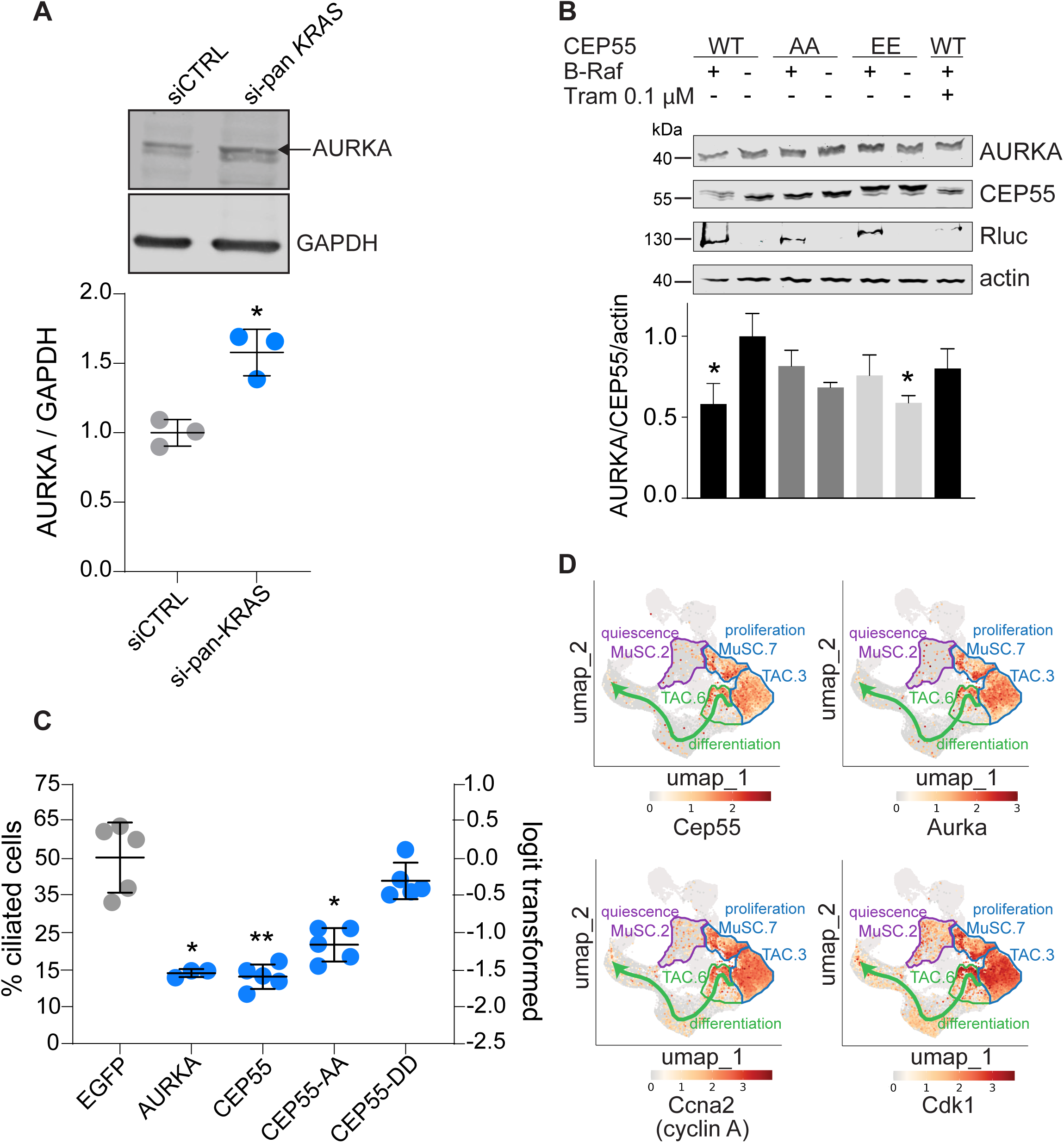
CEP55 and AURKA are highly expressed in cells transitioning from a ciliated progenitor-like to non-ciliated committed cells. (A) Immunoblot data with normalized AURKA levels from lysates of C2C12 cells grown in low serum for 72 h after transfection with 100 nM of the indicated siRNA, N = 3. Means ± SD are plotted. Statistical analysis was done using the Mann-Whitney test. (B) Immunoblot analysis of HEK293 cells transfected with Rluc8-tagged B-Raf and Flag-tagged CEP55, CEP55-AA or CEP55-EE. Shown are representative images of immunoblots labelled with AURKA, CEP55, Renilla luciferase or actin (loading control) antibodies. The graph shows the quantification of normalized AURKA levels, N = 4, except for trametinib (Tram) treatment where N = 3. Means ± SEM are plotted. Statistical analysis was done with one-way ANOVA and Dunnett’s multiple comparisons test. Shown are the statistical comparisons with the CEP55 WT control condition. (C) C2C12 cells were transfected with EGFP alone or mEGFP-AURKA, mEGFP-CEP55, and mEGFP-CEP55-DD. Cells were cultured in high serum for 48 h thereafter and ciliated fractions were quantified with confocal microscopy. N ≥ 3, 300 cells analyzed per condition. Means ± SD are plotted. Statistical analysis was done with one-way ANOVA with Browne-Forsythe correction and Dunnett’s T3 multiple comparisons test. (D) C2C12 cell differentiation scRNAseq derived uniform manifold approximation and projection (UMAP) of Cep55, Aurka, Ccna2 (Cyclin A2) and Cdk1 normalized gene expression (from high and low serum samples). Relevant cell states and the differentiation trajectory are indicated.

Expression of AURKA significantly decreased ciliation of C2C12 cells, which was mimicked by expression of CEP55-AA and wt CEP55, suggesting the latter is mostly present in the non-phosphorylated state (**Figure 2C**). In agreement with our proposed mechanism, expression of CEP55-DD retained ciliation (**Figure 2C**).

CEP55 and AURKA gene expression data from our C2C12 cell differentiation scRNAseq dataset were consistent with their role during de-/ re-ciliation processes of asymmetrically dividing MuSC and TAC. High expression of both genes is seen at the border of clusters MuSC.7/ MuSC.2, MuSC.7/ TAC.3 and TAC.3/ TAC.6, i.e. associated with cell states MuSC.7, TAC.3 and TAC.6 (**Figure 2D**), which we previously identified as bona fide ciliated (Chippalkatti *et al*., 2026). These borders also show a high expression of proliferation and G2/ M-phase markers *Cdk1* and cyclin A2 (*Ccna2*) (**Figure 2D**), consistent with asymmetrically dividing cells undergoing de-/ and re-ciliation cycles.

In conclusion, phosphorylation of CEP55 on S423/S426 downstream of K-Ras-MAPK-signaling significantly reduces its association with AURKA, which becomes destabilized and can thus no longer decrease ciliation. Given the localization of CEP55 on the basal body and the fact that CEP55-DD downmodulates the master regulator of de-ciliation, AURKA, we propose that ciliary K-Ras-MAPK signaling leads to the phosphorylation of CEP55 on the basal body to sustain ciliation.

### PDE6D-associated proteins regulate ciliation and C2C12 cell differentiation

Using mathematical modelling we have previously shown that ciliation directs asymmetric cell divisions of the proliferating MuSC and TAC populations of the C2C12 skeletal muscle cell line (**Figure 3A**). Given the significant impact of ciliary K-Ras on ciliation and the dependence of K-Ras localization to the cilium on the trafficking chaperone PDE6D, we hypothesized that the PDE6D-associated machinery affects ciliation and differentiation like K-Ras (**Figure 3B**). In agreement with a significant role of PDE6D for ciliation, PDE6D knockout MEF cells remained essentially non-ciliated (**Figure S1A**). Similarly, knockdown of PDE6D in C2C12 cells reduced ciliation significantly almost to the same extent as depletion of Ift88 (**Figure 3C; Figure S1B**) (Chippalkatti *et al*., 2026).

**Figure 3:**
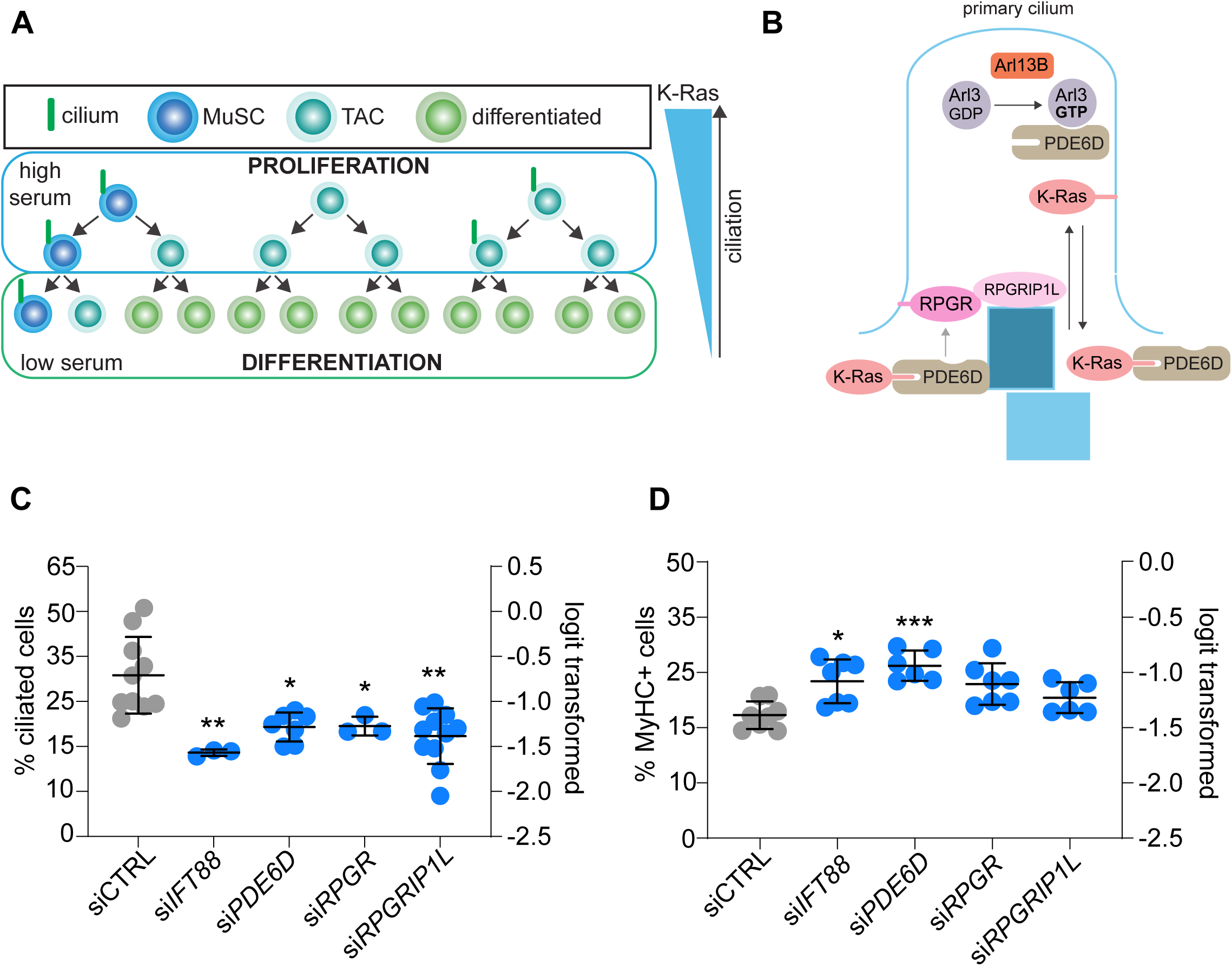
Proteins associated with PDE6D regulate ciliation and differentiation. (A) Schematic shows C2C12 cell populations and differentiation hierarchy with asymmetric cell divisions of MuSC being guided by the primary cilium. (B) Schematic of protein interactions associated with PDE6D in the cilium and on the basal body (dark blue rectangle). (C) Confocal imaging-based quantification of ciliation of C2C12 cells grown in low serum for 72 h after transfection with the indicated siRNAs (100 nM), N ≥ 3, n > 500 per condition. Means ± SD are plotted. Statistical analysis was done with one-way ANOVA with the Browne-Forsythe test and the Dunnett’s T3 multiple comparisons test. (D) Flow cytometric quantification of terminal differentiation marker MyHC expression in C2C12 grown in low serum for 72 h after transfection with the indicated siRNAs (100 nM), N ≥ 6. Means ± SD are plotted. Statistical analysis was done with one-way ANOVA with Browne-Forsythe correction and Dunnett’s T3 multiple comparisons test.

PDE6D possesses additional interaction partners, which could at least transiently dock it and potentially also its cargo such as K-Ras to the base of the cilium. RPGR localizes to the transition zone of cilia (Hong *et al*, 2003). It interacts via its N-terminal RCC1 domain with PDE6D in a manner that is compatible with the concurrent binding of PDE6D-cargo and release factors Arl2/3 (Watzlich *et al*, 2013). Both the ORF15 isoform of RPGR (RPGR-ORF15) and its interaction partner RPGRIP1L localize to centrioles, including the basal body of the primary cilium (Khanna *et al*, 2009; Shu *et al*, 2005; Vierkotten *et al*, 2007). Germline loss of RPGRIP1L is lethal and results in severe ciliopathy phenotypes (Delous *et al*, 2007). Similarly, germline loss-of-function mutations in PDE6D and RPGR are both associated with ciliopathies (Patnaik *et al*, 2015; Thomas *et al*, 2014), underscoring their significance during development.

We therefore tested if depletion of PDE6D-associated ciliary proteins RPGR and RPGRIP1L could also impact on ciliation and differentiation. Indeed, depletion of RPGR and RPGRIP1L significantly reduced ciliation to a similar extent as PDE6D depletion (**Figure 3C; Figure S1C,D**), while differentiation of C2C12 cells was increased with both manipulations, but not as significantly as after IFT88 or PDE6D depletion (**Figure 3D**).

Altogether these data support a substantial role of the PDE6D machinery in controlling ciliation and cell differentiation of C2C12 cells.

### Phosphomimetic mutation of the C-terminal Ser181 of KRAS reduces PDE6D-binding and ciliary abundance of KRAS

To understand how the PDE6D machinery could regulate ciliary abundance of K-Ras, we focused our attention on Ser181 at the C-terminus of K-Ras, which was proposed to reduce PDE6D affinity of K-Ras when phosphorylated (Dharmaiah *et al*., 2016).

We first explored mutations of this residue that would render it non-phosphorylatable or would mimic the negative charge of the phosphate. Intracellular PDE6D interaction studies using BRET confirmed a slightly reduced BRETtop interaction value for the S181A K-Ras mutant, and a much lower value for the phosphomimetic S181D mutant of K-Ras (**Figure 4A**). In line with these BRET data, K-Ras S181A accumulated to almost the same extent as wt K-Ras in cilia of C2C12 cells (**Figure 4B**), while ciliary localization of K-RasS181D was significantly reduced to background levels (**Figure 4C,D**).

**Figure 4:**
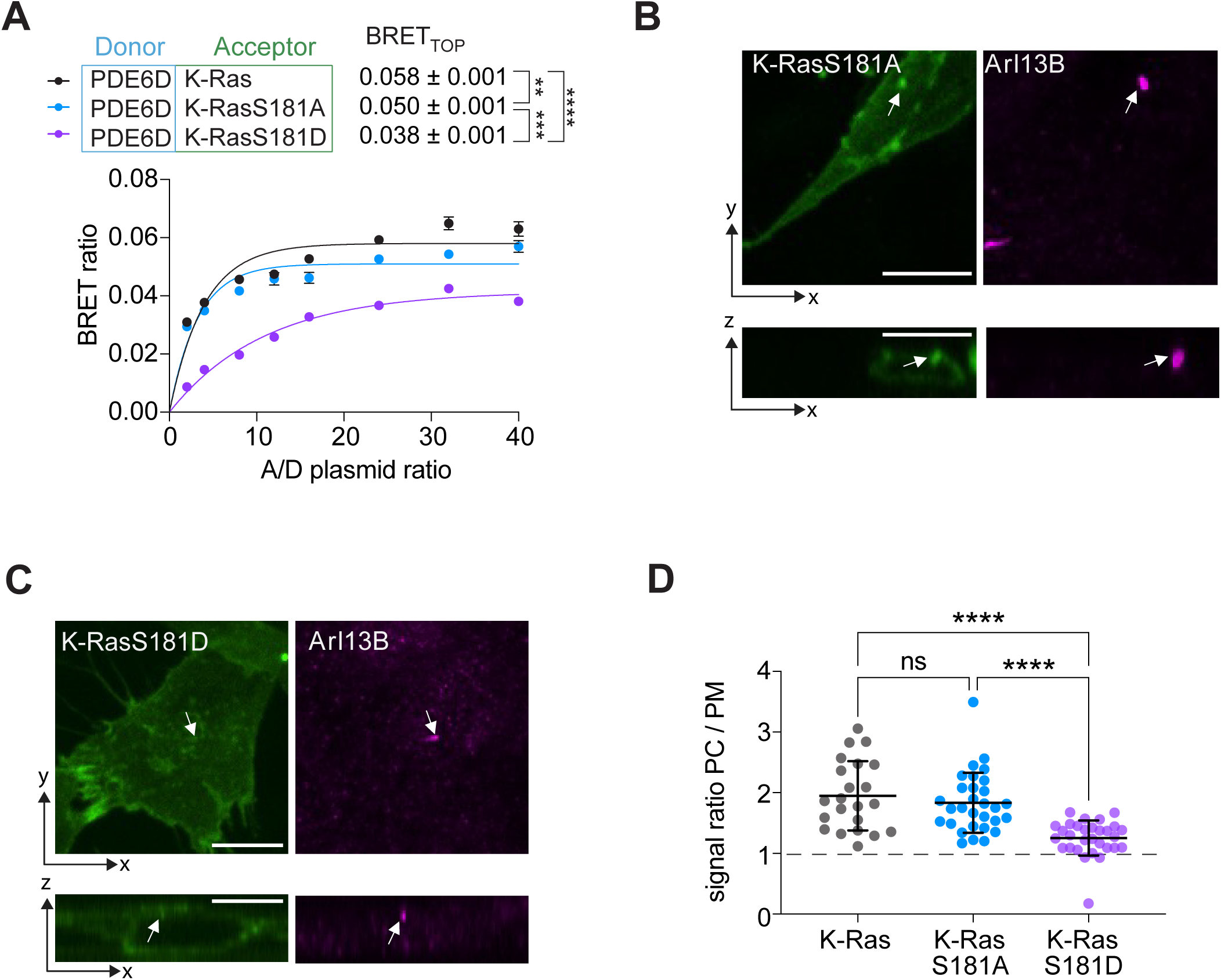
Modulation of phosphorylation status at S181 residue of K-Ras decreases BRET with PDE6D and reduces its ciliary accumulation. (**A**) BRET titration curves of Rluc8-PDE6D and GFP2-K-RasS181-mutants expressed in HEK293 cells, N = 3. Statistical analysis was performed with the extra sum-of-squares F-test. (**B, C**) Confocal images of C2C12 cells grown in high serum for 48 h after transfection with GFP2-K-RasS181A (B) or GFP2-K-RasS181D (C) and immunolabelled for ciliary marker Arl13B (arrows). *Bottom*, images show orthogonal views. Scale bar = 10 µm. (**D**) Quantification of Ras signal in the primary cilium (PC) as compared to the plasma membrane (PM) per cell using data as in (B,C), N = 3, GPF2-K-Ras, n = 22; GFP2-K-RasS181A, n = 30; GFP2-K-RasS181D, n = 30. Means ± SD are plotted. Statistical analysis was done with the Kruskal-Wallis test and Dunn’s post hoc test.

These results confirm that phosphorylation of Ser181 is sufficient to inhibit binding of K-Ras to PDE6D and demonstrate that this is sufficient to reduce the ciliary localization of K-Ras.

### Activation of PKG2 decreases PDE6D-binding and ciliary localization of KRAS

PKG2 was identified as a major kinase mediating phosphorylation of K-Ras on Ser181 (Cho *et al*, 2016). PKG2 is a cGMP-activated kinase downstream of the ciliary AMPK-pathway (Lai & Jiang, 2020) (**Figure 5A**).

**Figure 5:**
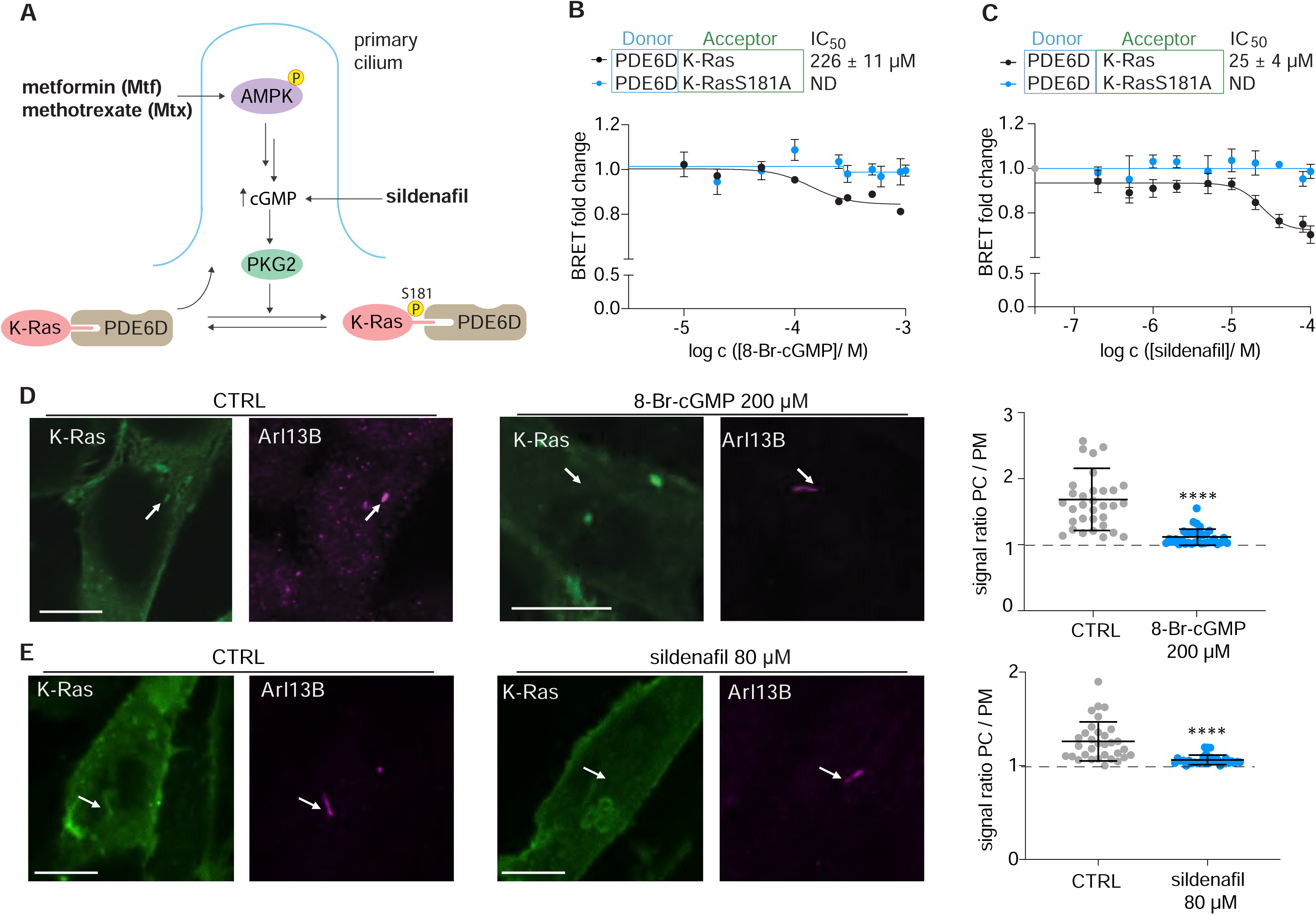
Pharmacological activation of the AMPK-PKG2 pathway disrupts K-Ras/ PDE6D interaction, thereby decreasing ciliary K-Ras localization. (A) Schematic showing how the AMPK-PKG2 pathway promotes phosphorylation of K-Ras on S181, which reduces binding to PDE6D and thus ciliary trafficking. Approved drugs modulating pathway activity are indicated in bold. (**B, C**) Dose-dependent effect of 8-Br-cGMP (B) or sildenafil (C) treatment for 24 h on BRET-interaction between Rluc8-PDE6D and GFP2-K-Ras or GFP2-K-Ras-S181A in HEK293 cells, N = 3. Donor-to-acceptor ratio was fixed at 1:8. (**D, E**) Confocal images of C2C12 cells expressing GFP2-K-Ras and treated as indicated in high serum for 48 h and immunolabelled for ciliary marker Arl13B (arrows). Scale bar = 10 µm. CTRL is water (D) or 0.1 % DMSO in high serum (E). Quantification of Ras signal in the primary cilium (PC) as compared to the plasma membrane (PM) per cell is shown to the right, N = 3, n = 33 (D); n = 31 (E). Means ± SD are plotted Statistical analysis was done with the Mann-Whitney test.

PKG2 activation can be stimulated by the analogue 8-Br-cGMP or clinically approved inhibitors of cGMP-degrading phosphodiesterase-5 (PDE5), such as sildenafil (Cho *et al*., 2016). Activation of PKG2 with these drugs is therefore expected to block binding of K-Ras to PDE6D and thus its trafficking to the cilium. Indeed, both 8-Br-cGMP (IC50 = 226 ± 11 μM) (**Figure 5B**) and sildenafil (IC50 = 25 ± 4 μM) (**Figure 5C**) dose-dependently decreased the BRET between K-Ras and PDE6D, while having no effect on non-phosphorylatable K-RasS181A. Accordingly, localization of K-Ras to the cilium was abrogated by treatment with 8-Br-cGMP (**Figure 5D**), while ciliary localization of K-RasS181A remained unaffected (**Figure S2A**). Likewise, a loss of ciliary K-Ras was observed after treatment with sildenafil (**Figure 5E**) or the related tadalafil (**Figure S2B**).

Hence, activation of the PKG2 increases Ser181-phosphorylation of K-Ras, which reduces its binding to PDE6D and thus ciliary localization of K-Ras.

### Stimulation of the AMPK-PKG2 pathway increases muscle cell differentiation

In line with the inverse correlation of ciliation and differentiation after K-Ras depletion (**Figure 3A**) (Chippalkatti *et al*., 2026), or PDE6D-machinery depletion (**Figure 3C,D**), sildenafil treatment significantly increased differentiation (**Figure 6A**), while knockdown of PKG2 had the opposite effect (**Figure 6B; Figure S3A**).

**Figure 6:**
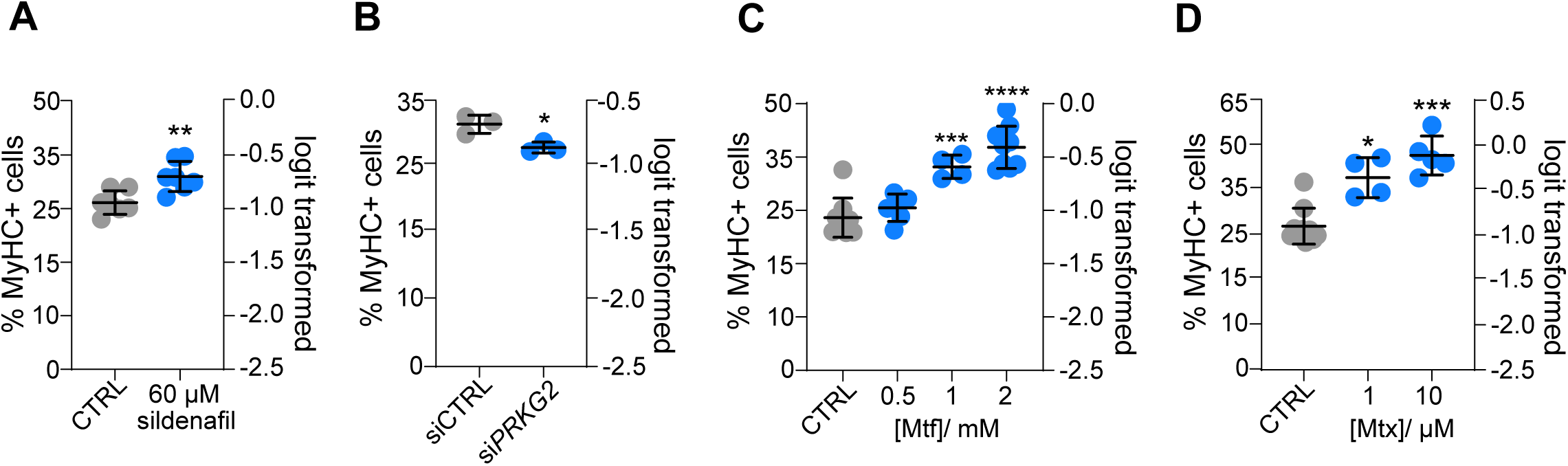
Activation of AMPK-PKG2 pathway through pharmacological agents increases C2C12 terminal differentiation. (A) Flow cytometric quantification of MyHC terminal differentiation marker expression in C2C12 cells treated as indicated for 72 h in low serum medium. CTRL (DMSO 0.1 % in low serum), N = 7. Means ± SD are plotted. Statistical analysis was done with one-way ANOVA with Browne-Forsythe correction and Dunnett’s T3 multiple comparisons test. (B) Flow cytometric quantification of MyHC terminal differentiation marker expression in C2C12 cells treated with PKG2 siRNA (100 nM) and cultured for 72 h in low serum. N = 3. Means ± SD are plotted. Statistical analysis was done with one-way ANOVA with Browne-Forsythe correction and Dunnett’s T3 multiple comparisons test. (**C, D**) Flow cytometric quantification of MyHC terminal differentiation marker expression in C2C12 cells treated as indicated for 72 h in low serum medium. CTRL (water) for metformin, Mtf (C), or CTRL (DMSO 0.1 % in low serum) for methotrexate, Mtx (D), N ≥ 4. Means ± SD are plotted. Statistical analysis was done with one-way ANOVA with Browne-Forsythe correction and Dunnett’s T3 multiple comparisons test.

PKG2 is positively regulated by the AMPK pathway, which can be activated by metformin or methotrexate (Beckers *et al*, 2006; Cho *et al*., 2016) (**Figure 5A**). Consistently, metformin and methotrexate dose-dependently increased differentiation (**Figure 6C,D**).

Analysis of the expression of the AMPK-PKG2-PDE6D machinery in our published scRNAseq dataset (Chippalkatti *et al*., 2026), revealed a broad expression of their genes across all muscle cell differentiation states (**Figure S3B-D**), where the highest expression, notably of PKG2, was observed in the quiescent MuSC.2 state. Together with the fact that this cell population has a low KRAS expression (Chippalkatti *et al*., 2026), we speculate that the high aggregate expression of the AMPK-PKG2 machinery indicates its increased employment. The resulting activity would keep the already low levels of K-Ras phosphorylated on Ser181 and thus out of the cilium to reduce ciliation, consistent with this cell state being non-ciliated (Chippalkatti *et al*., 2026).

## Discussion

The primary cilium has long been disregarded as a vestigial organelle, until the discovery of ciliopathies (Reiter & Leroux, 2017). Cyliary dysfunction can be expected to impact on key developmental processes, such as developmental signaling pathways, cell fate decisions and tissue patterning (Yamaguchi *et al*, 2025). However, our knowledge about how de- and re-ciliation are regulated originates from model systems where cilia are somewhat artificially induced, i.e. not integrated into their native, developmental context.

We recently described that K-Ras in the primary cilium of MuSC and TAC protects stemness or restricts commitment during asymmetric cell divisions, which can effectively reduce downstream differentiation (Chippalkatti *et al*., 2026). The predominant role of K-Ras is explained by the ciliary trafficking chaperone PDE6D, which prefers K-Ras over the other major Ras isoforms (Chandra *et al*., 2011; Dharmaiah *et al*., 2016; Siddiqui *et al*., 2020). The involvement of K-Ras in the regulation of ciliation could explain how it can broadly impact on developmental processes that depend on the primary cilium, such as asymmetric cell division. It is tempting to speculate that the more severe developmental phenotypes of K-Ras seen in knock-out mice is linked to this significant role during ciliation, as it is the sole and central mediator with little backup from the other major paralogs, N-Ras and H-Ras (Johnson *et al*, 1997).

Here we described two candidate pathways downstream of ciliary K-Ras that could regulate re-ciliation (K-Ras-CEP55-AURKA pathway) and de-ciliation (AMPK-PKG2-K-Ras pathway) at distinct stages of skeletal muscle cell proliferation and differentiation.

To support re-ciliation, we propose that ciliary K-Ras-MAPK activity promotes Ser425/428-phosphorylation of CEP55 to stabilize the ciliated states in proliferating MuSC and TAC during asymmetric divisions. Phospho-CEP55 binds less to AURKA, which reduces AURKA stability and thus counteracts its de-ciliation activity. AURKA is central to regulating de-ciliation, as it integrates various molecular inputs (Korobeynikov *et al*., 2017). The exact sequence of signaling events and their spatio-temporal context downstream of CEP55-AURKA that need to be enabled during asymmetric cell division remain at this point speculative. We propose that a high ciliary K-Ras-MAPK activity induces phosphorylation of CEP55 on the basal body, which essentially acts as a ‘stemness mark’ that may become asymmetrically apportioned during cell divisions of ciliated cells via the mother centrosome. Direct experimental validation is difficult, given the small population of ciliated cells and even more so the relatively small pool of ciliary K-Ras.

Others showed that the C-terminus of CEP55 is involved in stabilizing AURKA and thus disassembly of cilia but without considering any modulation of its activity by phosphorylation (Zhang *et al*., 2021). In line with their data, we also observed a loss of ciliation with the expression of wt CEP55 or CEP55-AA, while the phospho-mimetic CEP55-DD mutant protected ciliation. Furthermore, their in vitro data examined ciliation mostly in classical ciliogenesis models, such as serum starved MEF and RPE-1 cells, and not in the context of muscle cell differentiation.

At this point it is not clear whether phosphorylated CEP55 does not bind to AURKA due to electrostatic changes in its binding interface or due to its subcellular redistribution. Upon entry into mitosis, CEP55 becomes phosphorylated and dissociates from the centrosome, suggesting that subcellular redistribution could at least in part explain why phosphorylated CEP55 or its phospho-mimetic mutants do not bind to AURKA anymore. In line with this, both the MKS-mutant and a C-terminal ∼100 residue-deletion mutant do not localize to the centrosomes anymore (Fabbro *et al*., 2005; Zhang *et al*., 2021). Inheritance of phosphorylated CEP55 via the basal body-derived mother centrosome would in the daughter cells asymmetrically disable AURKA. This would permit re-ciliation in only the phospho-CEP55 inheriting daughter cell, which would retain stemness properties due to the re-establishment of ciliary stemness signaling pathways (Chen & Yamashita, 2021). However, observing such processes directly by time-lapse imaging is technically highly challenging.

Regarding the role of K-Ras during de-ciliation, we propose that the ciliary AMPK-PKG2-pathway activity increases K-Ras Ser181-phosphorylation, which reduces its binding to PDE6D and thus recruitment to the cilium. Hence this pathway would negatively impact on ciliation of TAC and most notably MuSC, where AMPK and PKG2 expression levels are elevated. The AMPK-PKG2 pathway may therefore serve as a physiological lever to regulate not only ciliation during cell differentiation but potentially other properties of cilia, such as length.

The elevated expression of the AMPK-PKG2-PDE6D machinery is seen in the committed and differentiating clusters TAC.1 and DIFF.0, which may suggest that their activity increase is required to promote complete de-ciliation and differentiation by preventing K-Ras from entering the cilium. During differentiation, myoblasts transiently still possess a cilium, similar to what we surmise for our TAC.6 population (Ng *et al*, 2021). It was proposed that cilia on differentiating myoblasts may serve a homing function to sites of injury. However, ultimately, muscle fibers lose cilia during terminal differentiation (Ng *et al*., 2021; Palla *et al*, 2022).

A particularly high AMPK-PKG2 gene expression is associated with the non-ciliated, quiescent MuSC state, implying a significance of the pathway to maintain a distinct quiescent state. In line with the general concept that quiescence induced by serum withdrawal induces ciliation, others suggested that quiescent MuSC remain ciliated (Jaafar Marican *et al*., 2016). However, multiple MuSC states exist in vivo that may be characterized by various depths of quiescence, with and without cilia (Barruet *et al*, 2020).

We therefore here propose that the bona fide non-ciliated MuSC.2 state corresponds to a deeper quiescence, associated with the induction of caveolin-1 and SPRY1 (Barruet *et al*., 2020; Chippalkatti *et al*., 2026). We recently showed that SPRY2, the paralog of SPRY1, directly binds to active K-Ras and blocks its access to effectors (Pavic *et al*, 2025). Hence, after SPRY1 levels decline a pool of active K-Ras could be released in MuSC.2 state cells, ready to restart the cell cycle. This would be in line with the observed increase in MAPK-activity upon exit from quiescence of MuSC (Machado *et al*, 2021). Once AMPK-activity is suppressed, such as after actual mitogen stimulation, K-Ras access to the cilium would allow for re-ciliation and transition into the adjacent MuSC.7 state (Chippalkatti *et al*., 2026).

Alternatively, the MuSC.2 state is an artificial aged state, as the loss of cilia with age on MuSC impairs injury-induced muscle repair in vivo (Palla *et al*., 2022). Indeed, the AMPK-pathway as a major integrator of environmental and intracellular cues, may continuously impact on ciliation efficiency as asymmetric cell divisions do not seem to progress with absolute efficiency, i.e. if insufficient active K-Ras is in the cilium. In such a case, a ciliated stem cell would not self-renew but instead give rise to two non-ciliated, committed TAC, leading to the long-term attrition of stem cells as observed during aging.

Yet, proper and timely re-activation of the AMPK-pathway may be crucial for inducing re-quiescence of stem cells after activation, which could otherwise suffer from the afore mentioned attrition process. Stem cell attrition may be particularly relevant in tissues such as muscle, which can undergo very dynamic anabolic and catabolic adaptation processes.

Our mechanisms implicate ciliary K-Ras activity with re- and de-ciliation processes in proliferating MuSC and TAC, and de-ciliation during MuSC quiescence and terminal differentiation, consistent with reports that implicated FGF-signaling in regulating cilia length and function (Kunova Bosakova *et al*., 2019). Given the persistent and significant role of FGF-signaling during mammalian development it is tempting to speculate that the most fundamental role of Ras during development is executed from the primary cilium (Chippalkatti & Abankwa, 2021; Duval *et al*, 2025).

## Limitations of the Study

Future studies could monitor directly the signaling from K-Ras4B via the MAPK-pathway to CEP55 in the cilium and at the basal body. By following the subsequent relocalization of CEP55 in asymmetrically dividing cells one could demonstrate how it then modulates AURKA stability inside the cell. This could resolve further details of whether and how CEP55 acts as a molecular stemness mark downstream of ciliary K-Ras and how it becomes asymmetrically apportioned as ciliated stem and progenitor cells divide.

Likewise, monitoring AMPK-activity directly in the cilium and identifying the molecular complexes that could target K-Ras to the basal body would further clarify the here proposed mechanisms. Ultimately, it will be important to demonstrate the broader involvement of the Ras-MAPK pathway in ciliogenesis during cell differentiation and development.

## Resource Availability

### Lead Contact

Further information and requests for resources and reagents should be directed to and will be fulfilled by the lead contact, Prof. Dr. Daniel Kwaku Abankwa.

### Materials Availability

Plasmids generated in this study will be available from the lead contact upon request.

### Data and Code Availability

The mouse C2C12 cell line single-cell RNA sequencing data described here are available at the ENA database under the accession reference PRJEB89780. Code used in this study is listed in the Key Resources Table and has been deposited in Gitlab at https://gitlab.com/uniluxembourg/fstm/dlsm/ccbdd/chippalkatti_2025, GitHub at https://github.com/sysbiolux/RasCilium and https://github.com/GesaHlzer/BachelorThesis.

## Acknowledgements

We thank Julien Aymon for his technical support. We are grateful to Professor Richard A. Kahn (Emory University School of Medicine, Atlanta, GA, USA) for providing the MEF PDE6D KO cell line. We also thank Dr. Rashi Halder (genomics platform, LCSB, University of Luxembourg) for running scRNAseq samples and Dr. Anthoula Gaigneaux (Bioinformatics Core, DHML, University of Luxembourg) for helping analyze scRNAseq results.

This work was supported by grants from the Luxembourg National Research Fund (FNR) grant C19/BM/13673303-PolaRAS2, INTER/FWO/23/18086068 molGluRAS2 and AFR/23/ 17112420/Bil ABANKWA SPREDCanUL2 to D.K.A.

## Author contributions

D.K.A conceived the project, supervised the study and acquired funding. R.C., E.S.R., B.P., C.L., A.G.-M. and Z.G., conducted experiments, analyzed and interpreted data and prepared figures. R.C., E.S.R. contributed to the drafting and D.K.A. wrote the final manuscript. All authors gave final approval and agreed to be accountable for all aspects of work, ensuring integrity and accuracy.

## Declaration of Interests

The authors declare no competing interests.

## Supplemental Information

Supplemental information can be found online at [link]

Document S1. Figures S1–S3

## Materials and Methods

### KEY RESOURCES TABLE

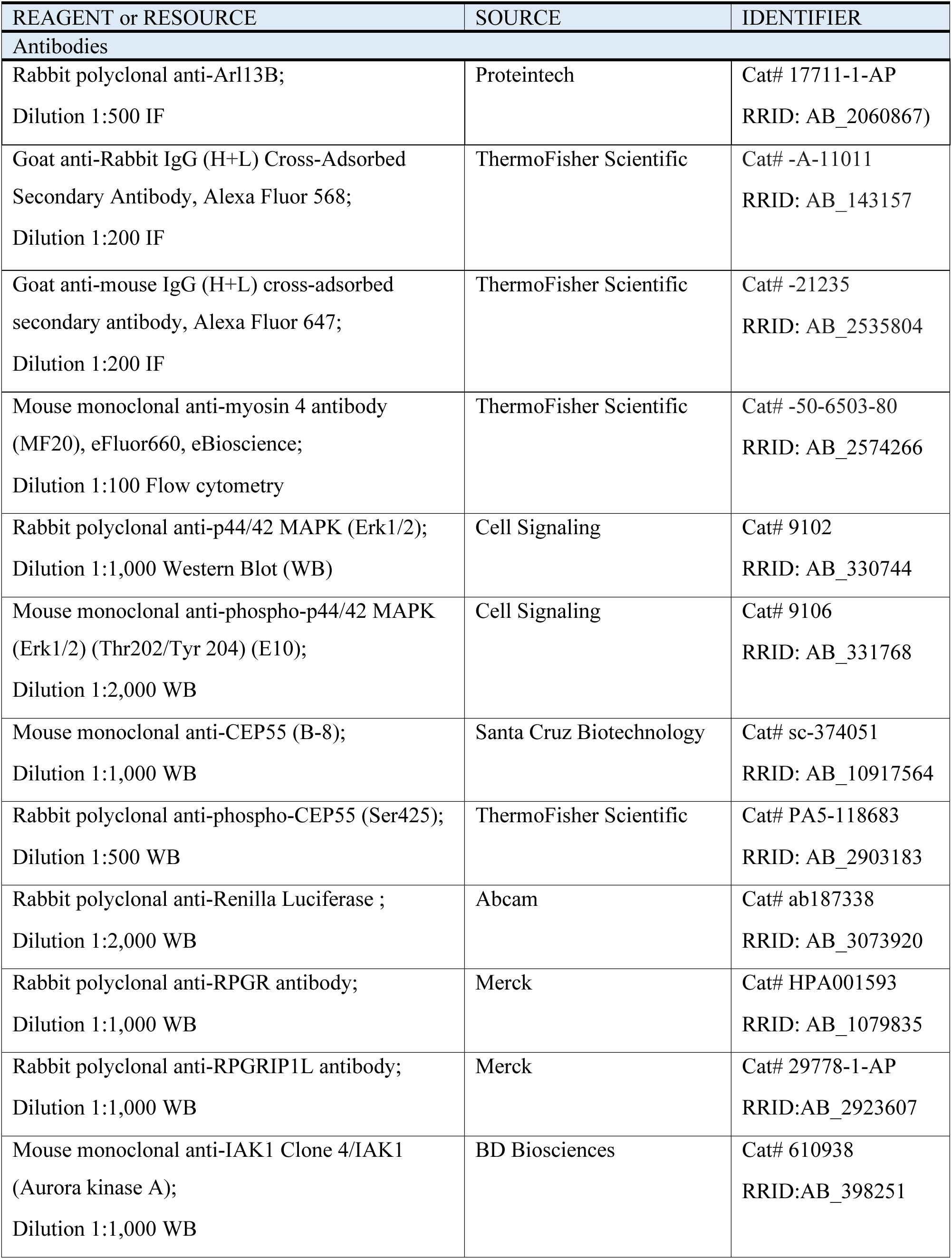
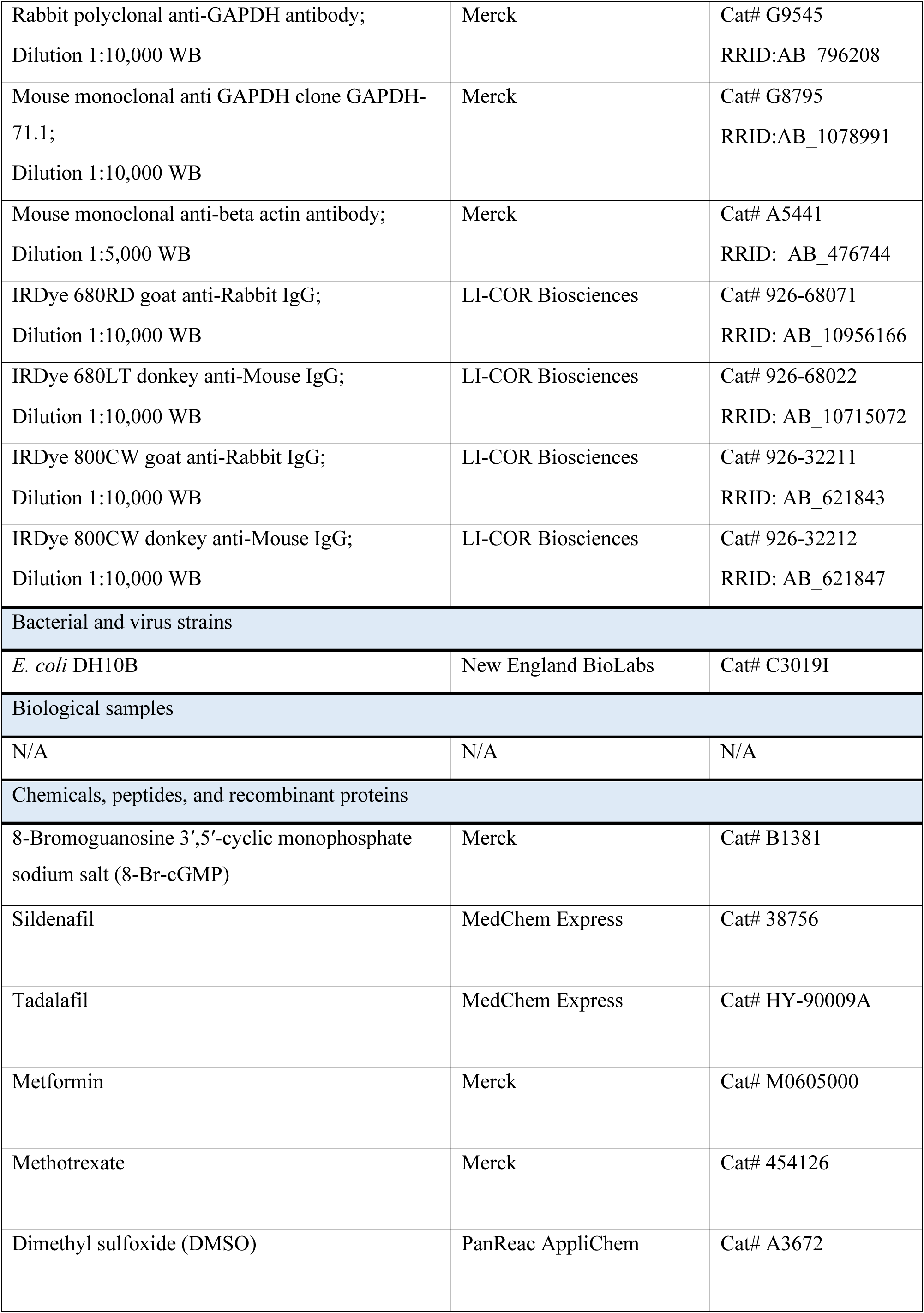
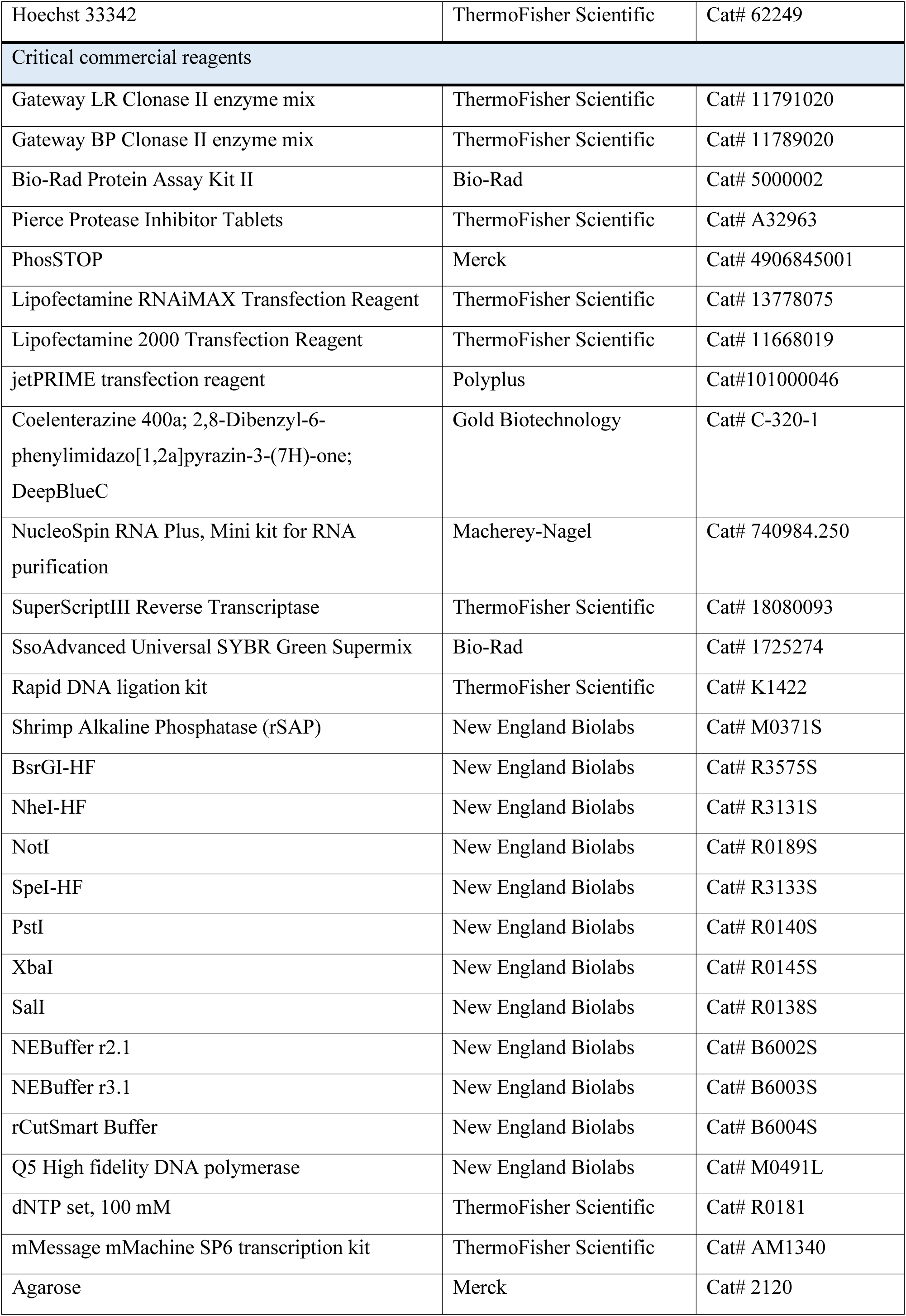
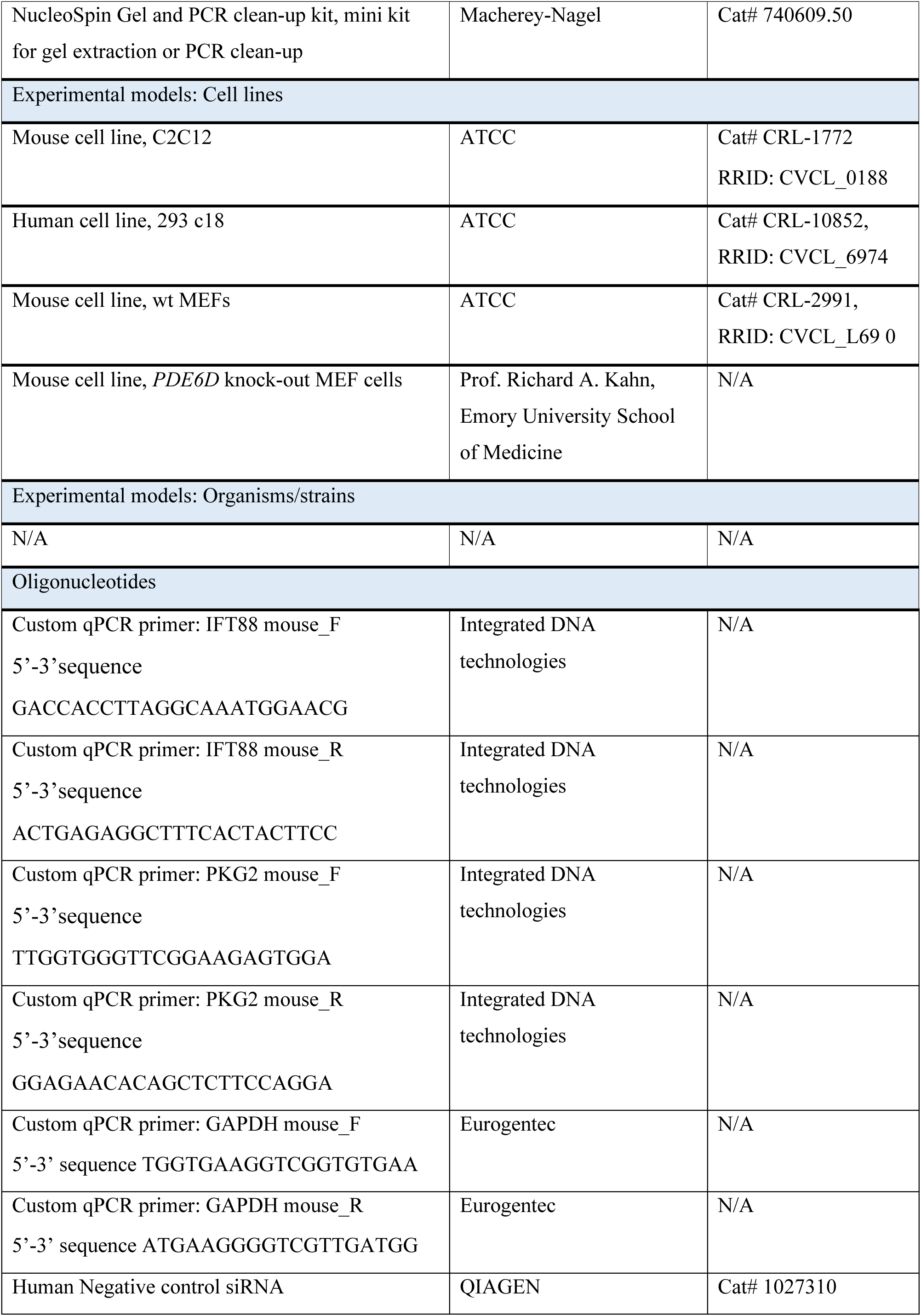
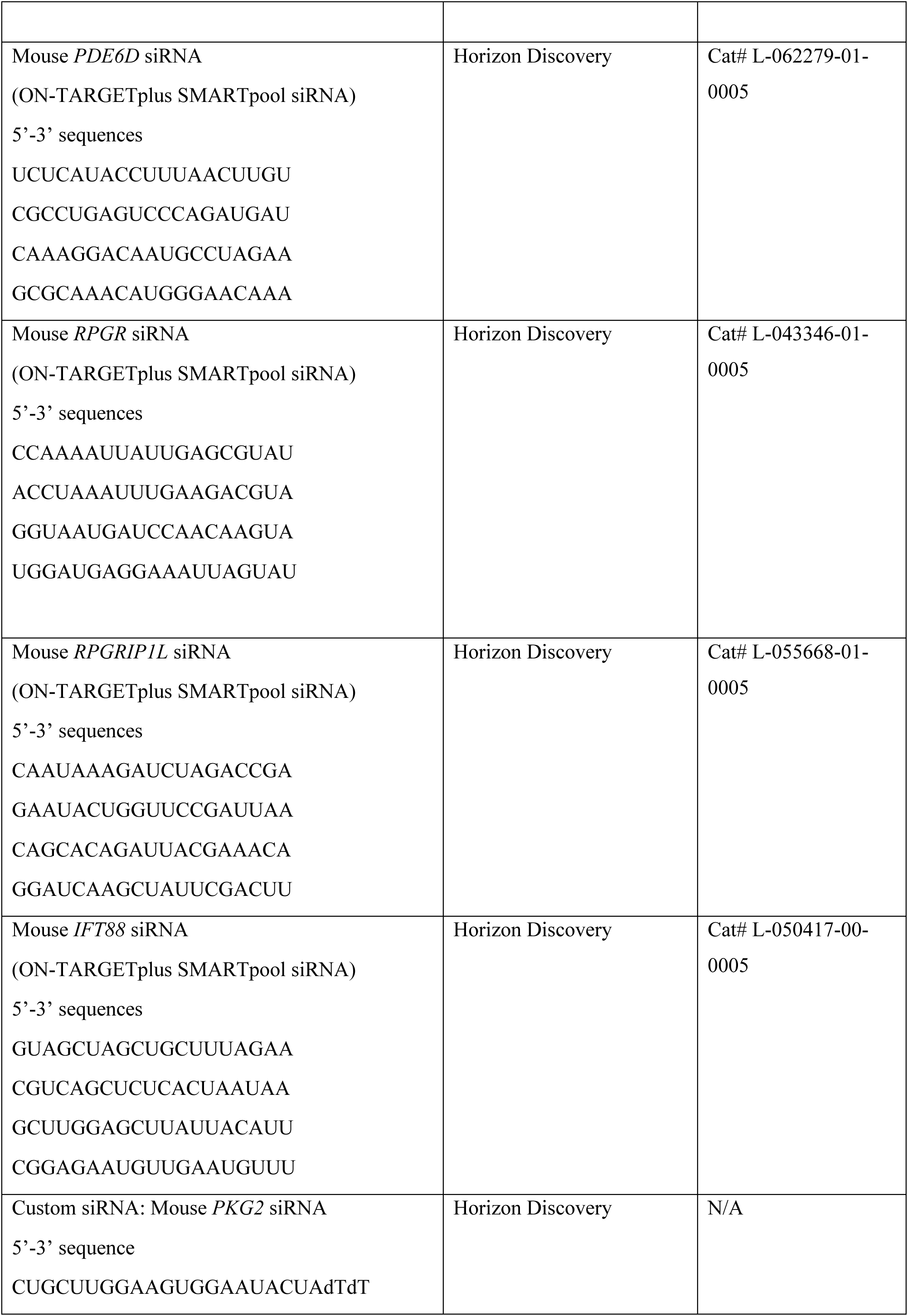
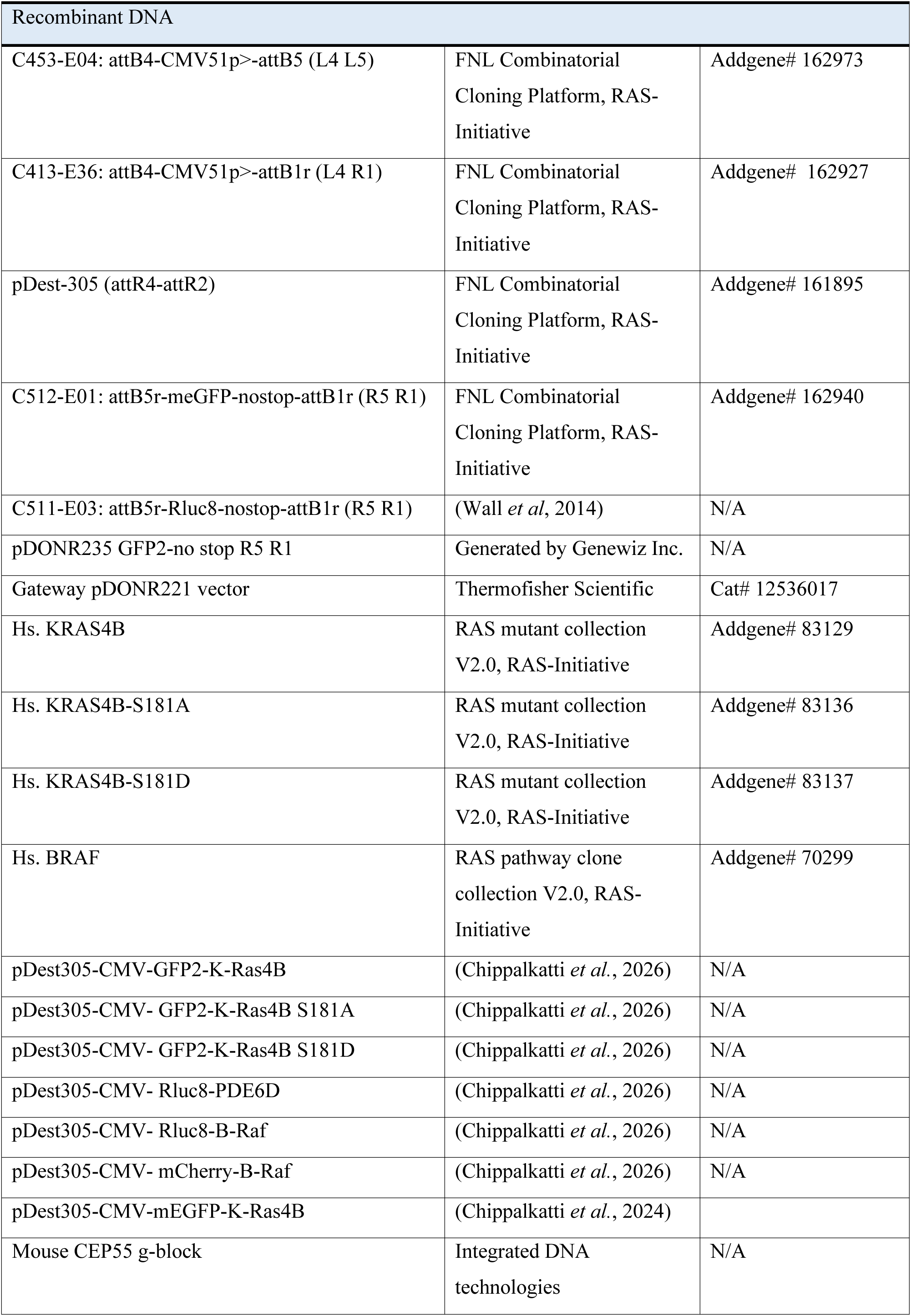
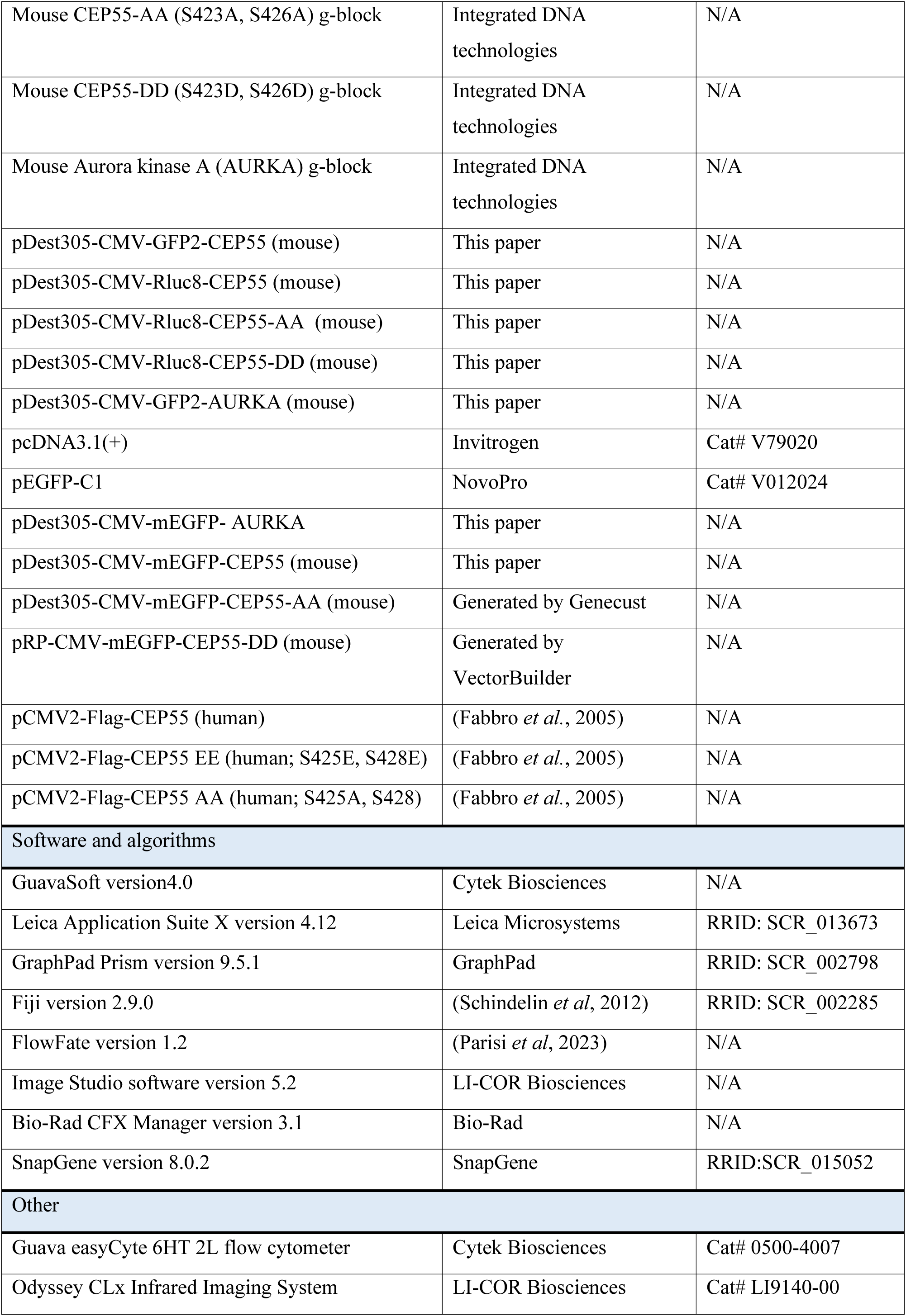
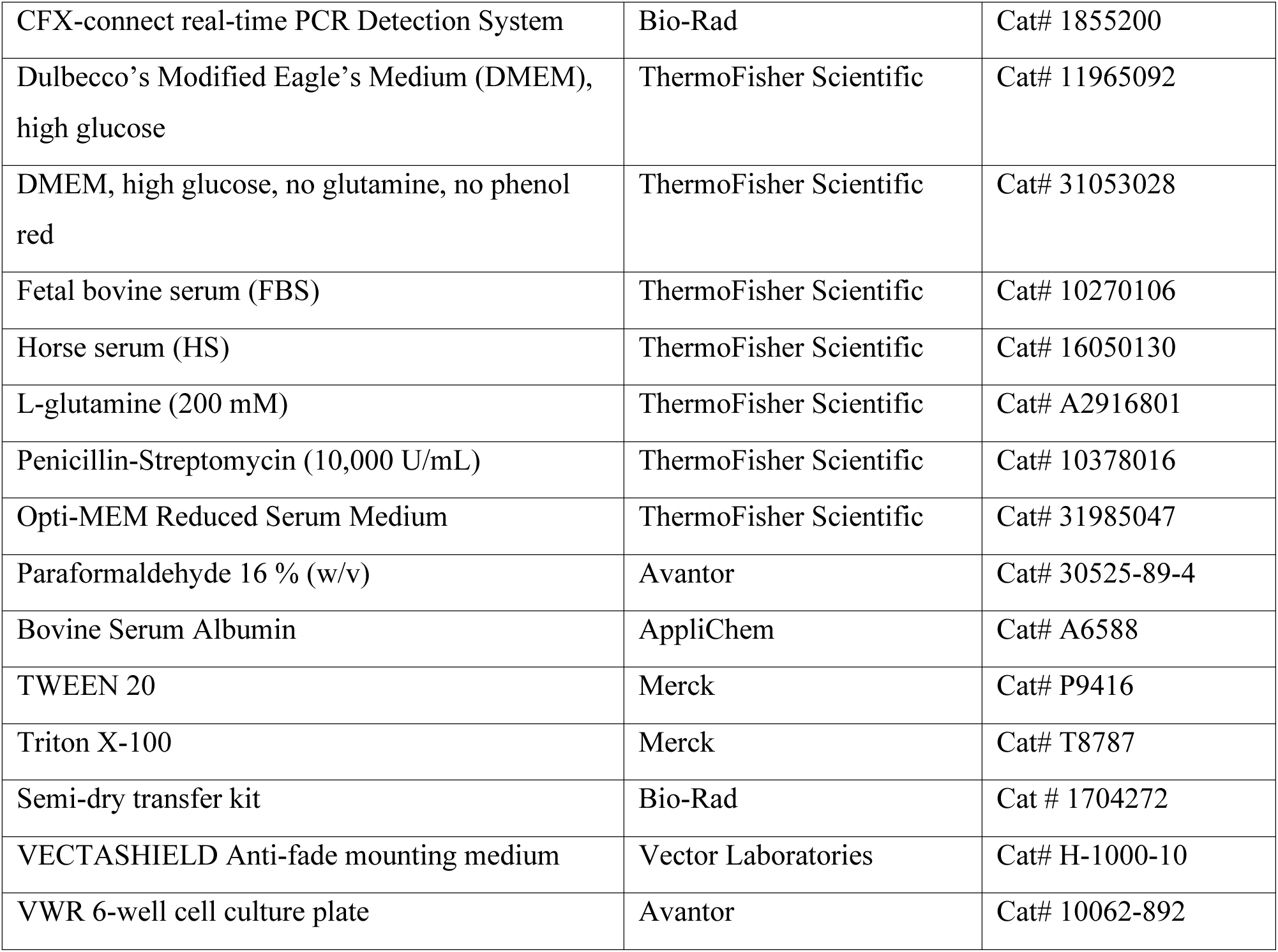

## EXPERIMENTAL MODEL AND STUDY PARTICIPANT DETAILS

### Cell lines

Cells were cultured in Dulbecco’s Modified Eagle’s Medium (DMEM) containing approximately 9% (v/v) fetal bovine serum (FBS), 2 mM L-glutamine, and 1% penicillin–streptomycin. For C2C12 cell culture experiments, this formulation is hereafter referred to as the “high-serum medium.” Cultures were maintained at 37 °C in a humidified atmosphere with 5% CO₂ and were routinely passaged twice weekly upon reaching 50–60% confluency. C2C12 differentiation experiments were performed as previously reported (Chippalkatti *et al*., 2024; Parisi *et al*., 2023). In brief, cells were grown to approximately 90% confluency before induction of differentiation. Differentiation was initiated by replacing the high-serum medium with a low-serum medium consisting of DMEM supplemented with 2% horse serum. The low-serum medium was replenished daily for a period of three days. Details of the treatment conditions used in individual experiments are provided in the corresponding figure legends.

## METHOD DETAILS

### Plasmid construct generation

Expression constructs encoding B-Raf, CEP55 mutants, and PDE6D variants were generated using a MultiSite Gateway cloning strategy. For gene variants available as gBlock DNA fragments, entry clones were first created by recombining the gBlocks with the pDONR221 donor vector using BP Clonase II enzyme mix, thereby introducing the appropriate recombination sites required for subsequent LR reactions.

The resulting variant entry clones were then combined with entry clones carrying the CMV promoter and either Rluc8, GFP2, or mEGFP coding sequences, and recombined into the pDest305 or pDest312 destination vectors as previously described (Wall *et al*., 2014). LR Gateway recombination was performed using LR Clonase II enzyme mix, after which the reaction products were transformed into the ccdB-sensitive *Escherichia coli* strain DH10B. Ampicillin-resistant transformants were selected and screened by BsrGI-HF restriction enzyme digestion to verify the presence of the desired expression constructs. The integrity of all final plasmids was subsequently confirmed by Sanger sequencing.

### Cell transfection

C2C12 cells were transfected with the plasmid constructs specified in the corresponding figure legends using Lipofectamine 2000. Transfections were performed in cells maintained in high-serum medium at approximately 50–60% confluency. Plasmid DNA and transfection reagent quantities were used according to the manufacturer’s recommendations.

For each transfection, 2 µg plasmid DNA was diluted in 125 µL Opti-MEM and briefly mixed by vortexing, followed by a 5 min incubation at room temperature (23–25 °C). In parallel, 7 µL Lipofectamine 2000 was diluted in 125 µL Opti-MEM, vortexed briefly, and incubated for 5 min at room temperature. The diluted Lipofectamine 2000 solution was then combined with the diluted DNA, mixed gently, and allowed to form DNA–lipid complexes for 10 min at 23–25 °C.

The resulting transfection mixture was added dropwise to a single well of a 6-well plate containing cells in 2 mL of high-serum medium. After 4 h of incubation, the transfection medium was replaced with fresh high-serum medium. For differentiation experiments, cells were switched to low-serum medium 24 h post-transfection, and the medium was subsequently renewed daily for three consecutive days.

### siRNA transfection

siRNA-mediated knockdown experiments were carried out using Lipofectamine RNAiMAX reagent. C2C12 cells maintained in high-serum medium were transfected upon reaching approximately 50–60% confluency. For transfections performed in 6-well plates, siRNA was typically used at a final concentration of 100 nM and prepared in 250 µL Opti-MEM containing 7.5 µL Lipofectamine RNAiMAX.

The siRNA–lipid complexes were added directly to wells containing 2 mL of high-serum medium. After 24 h, the culture medium was exchanged for low-serum medium to initiate differentiation. The low-serum medium was subsequently replenished every 24 h for a total of three days.

### Immunofluorescence

For immunofluorescence analyses, C2C12 cells were seeded onto sterile 0.17 mm-thick glass coverslips in 6-well plates at a density of 100,000 cells/ mL. 24 h after seeding, cells were transfected with the fluorescent protein-tagged constructs indicated in the corresponding figure legends. Following transfection, cells were maintained either in high-serum medium for 48 h or in low-serum medium for 72 h, as specified for each experiment.

For experiments involving AMPK–PKG2 modulators, compounds were diluted in high-serum medium to the indicated final concentrations and added to the cultures 24 h after transfection. Cells were then incubated with the treatments for an additional 48 h.

Cells were fixed with 4% (w/v) paraformaldehyde in PBS and permeabilized using 0.5% Triton X-100 for 10 min. Following permeabilization, cells were washed with PBS containing 0.05% Tween 20 (PBST) and blocked for 30 min in 2% BSA prepared in PBST. Primary antibody incubations were carried out for 1 h, after which cells were washed three times for 5 min each with PBST. Samples were then incubated for 1 h with the appropriate secondary antibodies. After a further 5 min PBST wash, nuclei were stained with Hoechst 33342 (0.2 µg/mL in PBST) and washed once more with PBST.

Coverslips were mounted onto glass microscope slides using Vectashield mounting medium and prepared for imaging.

### Confocal imaging and quantification of ciliation

Fixed samples were imaged using a Nikon Ti-E microscope equipped with a Yokogawa CSU-W1 spinning-disk confocal unit and an Andor iXon Ultra EMCCD camera, using a 60× oil immersion objective (NA 1.3). GFP2- and mEGFP-tagged proteins were excited with a 488 nm laser and detected using EGFP emission settings with a 535/20 band-pass filter. Alexa Fluor 568 was excited using a 561 nm laser line and detected with a 560/40 band-pass filter, whereas Alexa Fluor 647 fluorescence was excited at 640 nm and detected with a 700/35 band-pass filter. Z-stack images were acquired at 0.3 μm intervals between optical sections. Image acquisition was performed using Nikon NIS-Elements software, and subsequent image analysis was carried out in Fiji/ImageJ (Schindelin *et al*., 2012).

To assess the relative enrichment of Ras constructs within primary cilia, three-dimensional orthogonal projections were generated in ImageJ/Fiji. Ciliary fluorescence intensity was determined by measuring the mean signal of GFP2- or mEGFP-tagged Ras constructs within a region of interest encompassing the primary cilium, identified by Arl13B staining. Plasma membrane fluorescence was measured as the mean signal intensity within a region of interest outlining the entire cell perimeter. Relative ciliary localization was expressed as the ratio of primary cilium to plasma membrane fluorescence intensity (PC/PM).

For quantification of the effect of mEGFP-tagged AURKA and CEP55 mutants on ciliation, a total of 100 mEGFP positive cells were counted at random per condition, per technical repeat and scored for the presence or absence of the cilium, identified by anti-Arl13B immunostaining. This experiment was repeated three times and a total of 300 cells per condition were quantified this way. The average percentage obtained across these images was reported as the percentage of ciliated cells.

### Flow cytometry-based C2C12 cell differentiation assay

Flow cytometric assessment of C2C12 differentiation was performed using a Guava easyCyte HT-2L flow cytometer, as described previously (Chippalkatti *et al*., 2024; Parisi *et al*., 2023). For each experimental repeat, more than 1,000 cells were analyzed within the final target gate. Briefly, C2C12 cells were seeded in 6-well plates at a density of 100,000 cells/ mL and subjected to the treatments indicated in the corresponding figure legends. During differentiation, low-serum medium containing the indicated compounds was refreshed daily for three consecutive days. Cells were subsequently harvested by trypsinization for 5 min and collected by centrifugation at 500 × *g* for 5 min. Cell pellets were fixed in 4% (w/v) paraformaldehyde prepared in PBS for 10 min, washed with PBS, and permeabilized with 0.5% Triton X-100 in PBS for a further 10 min.

Following permeabilization, cells were washed with PBST and stained for 1 h at 4 °C using eFluor 660-conjugated anti-myosin 4 MF20 antibody (1:100 dilution in PBST), which recognizes myosin heavy chain (MyHC; Myh1 gene product). After staining, cells were pelleted by centrifugation at 500 × *g* for 5 min and resuspended in PBS prior to flow cytometric acquisition. Instrument settings and gating strategies were established using unstained controls and MyHC-positive controls, as described previously (Parisi *et al*., 2023). Analysis was restricted to intact cells, and differentiation was assessed by MyHC positivity, which was detected using 640 nm excitation and the Red-R (664/20) emission filter. The extent of differentiation was quantified as the percentage of MyHC-positive cells. Data analysis and quantification were performed using FlowFate.

### BRET assay

For BRET assays, 200,000 HEK293 c18 cells were seeded in 12-well cell culture plates. After 24 h, cells were transfected using 3 μL of jetPRIME transfection reagent, a donor construct tagged with Rluc8 and an acceptor construct tagged with GFP2 as previously described (Babu Manoharan *et al*, 2023; Duval *et al*, 2024; Okutachi *et al*, 2021).

For BRET-titration experiments for donor saturation, a constant 25 ng concentration of donor plasmid and an increasing concentration of acceptor plasmid (from 0 to 1,000 ng) were transfected. Empty pcDNA3.1 vector was added to reach an identical 1,050 ng total DNA load per well. After 48 h, BRET measurements were taken at 25 °C using a CLARIOstar plate reader and the BRET ratio was calculated as previously described (Duval *et al*., 2024). Briefly, the luminophore-specific signal channels were λex = 405 ± 10 nm and λem = 515 ± 10 nm for the GFP2-acceptor signal and, after addition of 10 μM coelenterazine 400a, λem = 410 ± 40 nm for the Rluc8-donor signal and λ = 515 ± 15 nm for the BRET signal. The BRET ratio was then plotted as a function of the acceptor/donor plasmid ratio. Data typically from three biological repeats, each acquired as quadruplicate technical replicates, were averaged and analyzed together using nonlinear regression in GraphPad Prism, applying a one-phase association equation. The resulting Y_MAX_ parameter, which represents the upper asymptote of the fitted curve on the Y-axis, was taken as the BRET_TOP_ value, reflecting the maximal BRET ratio achievable within the tested acceptor-to-donor range. When the dataset did not exhibit a clear plateau, curve fitting was not applied; instead, the highest BRET ratio observed at the maximum acceptor/donor ratio was reported as the BRET_TOP_ value.

Dose–response BRET assays were performed following determination of the optimal acceptor-to-donor plasmid ratio by titration experiments, which are indicated in the respective figure legends. 24 h post-transfection, the culture medium was replaced with fresh medium containing increasing concentrations of the test compounds. Sildenafil was applied at 200 nM–100 µM, 8Br-cGMP at 10 µM–1000 µM and a 0.1% (v/v) DMSO solution in culture medium served as vehicle control. After an additional 24 h incubation, cells were harvested and processed for the BRET assay. For analysis, the logarithm of the inhibitor concentration was plotted against the BRET ratio, and data were fitted using the log (inhibitor) vs. response variable slope (four parameters) equation in GraphPad Prism to derive IC_50_ values.

### Immunoblotting

For cell transfection and subsequent lysis, 250,000 HEK293 c18 cells were seeded in 12-well plates. After 24 h, cells were transfected in Opti-MEM medium using 1 µg of plasmid DNA and 3 µL Lipofectamine 2000.

24 h post-transfection, cell lysis was performed by scraping cells in situ using ice-cold lysis buffer (50 mM Tris-HCl pH 7.5, 150 mM NaCl, 0.1 % w/v SDS, 5 mM EDTA, 1 % v/v Nonidet P-40, 1 % v/v Triton X-100, 1 % w/v sodium-deoxycholate, 1 mM Na_3_VO_4_, 10 mM NaF, 100 μM leupeptin and 100 μM E64D protease inhibitor) supplemented with protease and phosphatase inhibitor cocktails as previously described (Kaya *et al*., 2024). Lysates were clarified by centrifugation, and a Bradford assay was performed to determine total protein concentrations using a bovine serum albumin standard curve.

For SDS-PAGE, 40 μg of protein per lane were resolved under reducing conditions on 10 % w/v polyacrylamide gels and transferred to nitrocellulose membranes by semi-dry transfer. Following a saturation step of 1 h at room temperature in PBS 2 % w/v BSA 0.2 % TWEEN 20, membranes were incubated overnight at 4 °C with primary antibodies diluted in saturation buffer. For phospho-ERK or phospho-CEP55 quantification, combinations of mouse anti-phospho-ERK and rabbit anti-ERK or rabbit anti-phospho-CEP55 and mouse anti-CEP55 were used, respectively. Rluc8-tagged B-Raf was revealed with a rabbit anti-Renilla Luciferase antibody and Aurora kinase A with a mouse anti-IAK1 antibody. Actin was probed with a mouse anti-actin antibody as a loading control. After at least three wash steps in PBS 0.2 % v/v TWEEN 20, secondary antibody labeling was carried out for 1 h at room temperature, followed by three additional washes. Signal intensities were quantified using an Odyssey Infrared Image System (LI-COR Biosciences). For each condition, phospho-ERK/ERK and phospho-CEP55/CEP55 ratios were calculated. Aurora kinase A levels were normalized to CEP55 and actin. To account for technical variability between blots, ratios were normalized to the sum of all ratios within each blot. Data were then expressed relative to the averaged control condition and represented as mean ± SEM from at least N = 3 independent biological repeats.

### Quantitative RT-PCR

To validate siRNA-mediated gene silencing, C2C12 cells were seeded in 6-well plates at a density of 100,000 cells/ mL and transfected 24 h later with 100 nM negative control siRNA or siRNAs targeting mouse *PRKG2* (NM_008926.4) and *IFT88* (NM_009376.3). Following 72 h of culture under low-serum differentiation conditions, cells were harvested for total RNA isolation.

Total RNA was extracted using the NucleoSpin RNA Plus Mini kit according to the manufacturer’s protocol. Reverse transcription was performed on 1 µg of total RNA using SuperScript III Reverse Transcriptase to generate cDNA. Quantitative PCR analysis was carried out using SsoAdvanced Universal SYBR Green Supermix on a CFX Connect Real-Time PCR System (Bio-Rad), and data acquisition and analysis were performed using Bio-Rad CFX Manager Software.

Primer pairs for *IFT88*, *PRKG2* (PKG2), and *GAPDH* amplification were designed using the OligoPerfect Primer Designer online tool based on the corresponding mouse mRNA reference sequences: *IFT88* (NM_009376.3), *PRKG2* (NM_008926.4), and *GAPDH* (NM_008084.4). Relative transcript abundance was determined using the 2^−ΔΔCt method, with *GAPDH* serving as the reference gene for normalization.

## QUANTIFICATION AND STATISTICAL ANALYSIS

Unless otherwise indicated, data are presented as mean ± standard deviation (SD). Statistical analyses and graph generation were performed using GraphPad Prism version 9 or 10. The number of independent biological repeats (N) and individual data points (n) analyzed for each experiment are provided in the corresponding figure legends.

Representative microscopy images depict phenotypes observed across at least three independent biological repeats (N ≥ 3). For datasets expressed as percentages of cells, proportions were converted to continuous values by logit transformation of the cell fraction (*p*) according to: logit(p) = ln[p/(1 − p)]

Plots of logit-transformed data were displayed with the transformed scale on the right-hand Y-axis and the corresponding approximate percentage values on the left-hand Y-axis to facilitate interpretation. The statistical tests used for individual datasets are specified in the relevant figure legends. Unless otherwise stated, comparisons were performed relative to the designated control group, which is typically shown in grey. Statistical significance was defined as P < 0.05 and denoted as follows: ns, not significant; P < 0.05 (*); P < 0.01 (**); P < 0.001 (***); and P < 0.0001 (****).

## Notes

### Competing Interest Statement

The authors have declared no competing interest.

