## Supplementary Information for "Regulation of de- and reciliation by KRAS during muscle cell differentiation"

### Supplementary Figures

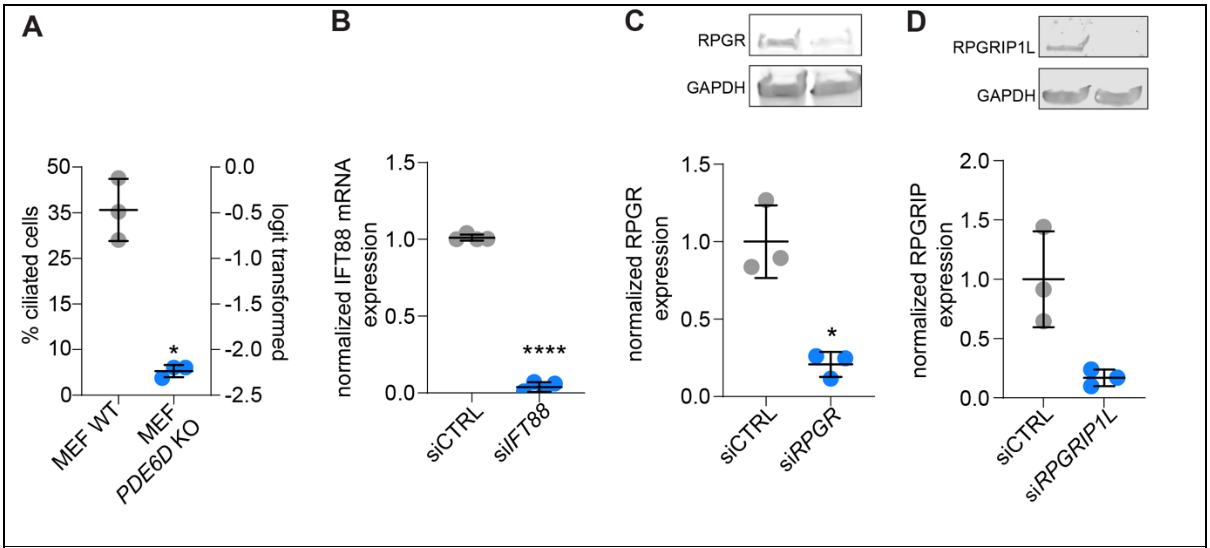


**Figure S1, related to Figure 3.**

(**A**) Confocal imaging-based quantification of ciliation of MEF wt and MEF PDE6D knockout (KO) cells after 24 h culture period in serum-free medium, N = 3, n > 600 per condition. Means ± SD are plotted. Statistical analysis was done with the unpaired t-test with Welch's correction.

(**B**) RT-PCR based quantification of *IFT88* mRNA normalized to *GAPDH* mRNA of C2C12 cells grown in low serum for 72 h after transfection with siCTRL or si*IFT88* (each 100 nM), N=2. Means ± SD are plotted. Statistical analysis was done with the unpaired t-test with Welch's correction.

(**C,D**) Representative immunoblots and quantification of protein levels normalized to GAPDH of C2C12 cells grown in low serum for 72 h after transfection with siCTRL or with si*RPGR* (C), si*RPGRIP1L* (D), at 100 nM. N = 3. Means ± SD are plotted. Statistical analysis was done with the unpaired t-test with Welch's correction.

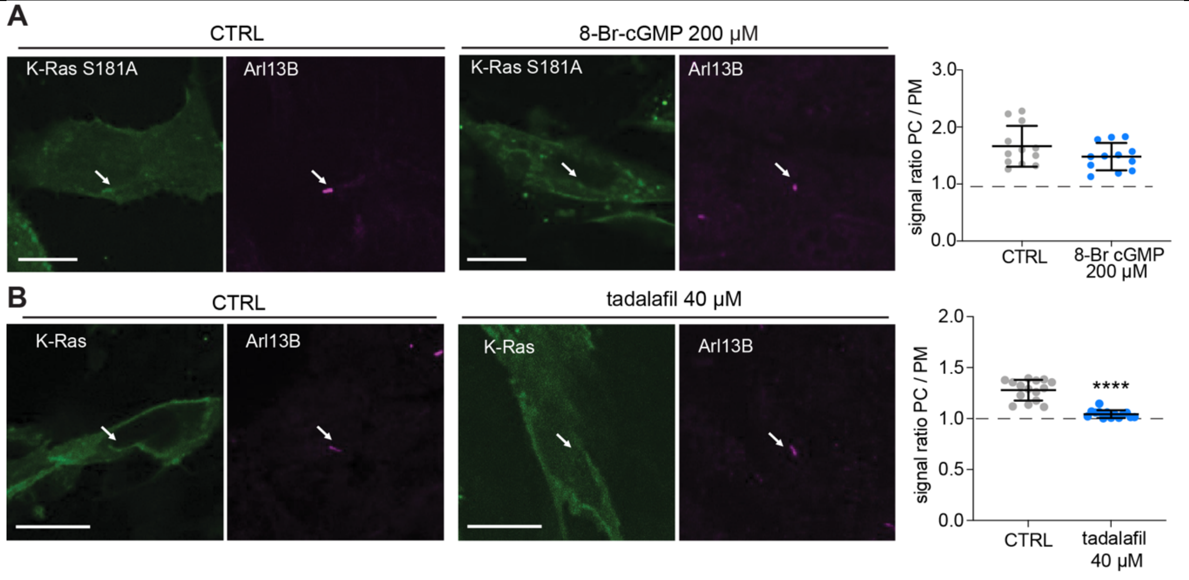


**Figure S2 related to Figure 5.**

(**A** ,**B**) Confocal images of C2C12 cells expressing GFP2-K-RasS181A (A) or GFP2-K-Ras (B) and treated as indicated in high serum for 48 h and immunolabelled for ciliary marker Arl13B (arrows). Scale bar = 10 µm. CTRL is water (A) or 0.1 % DMSO in high serum (B).

Quantification of K-Ras signal in the primary cilium (PC) as compared to the plasma membrane (PM) per cell is shown to the right, N = 3, n = 12 (A); n = 15 (B). Statistical analysis was done with the Mann-Whitney test.

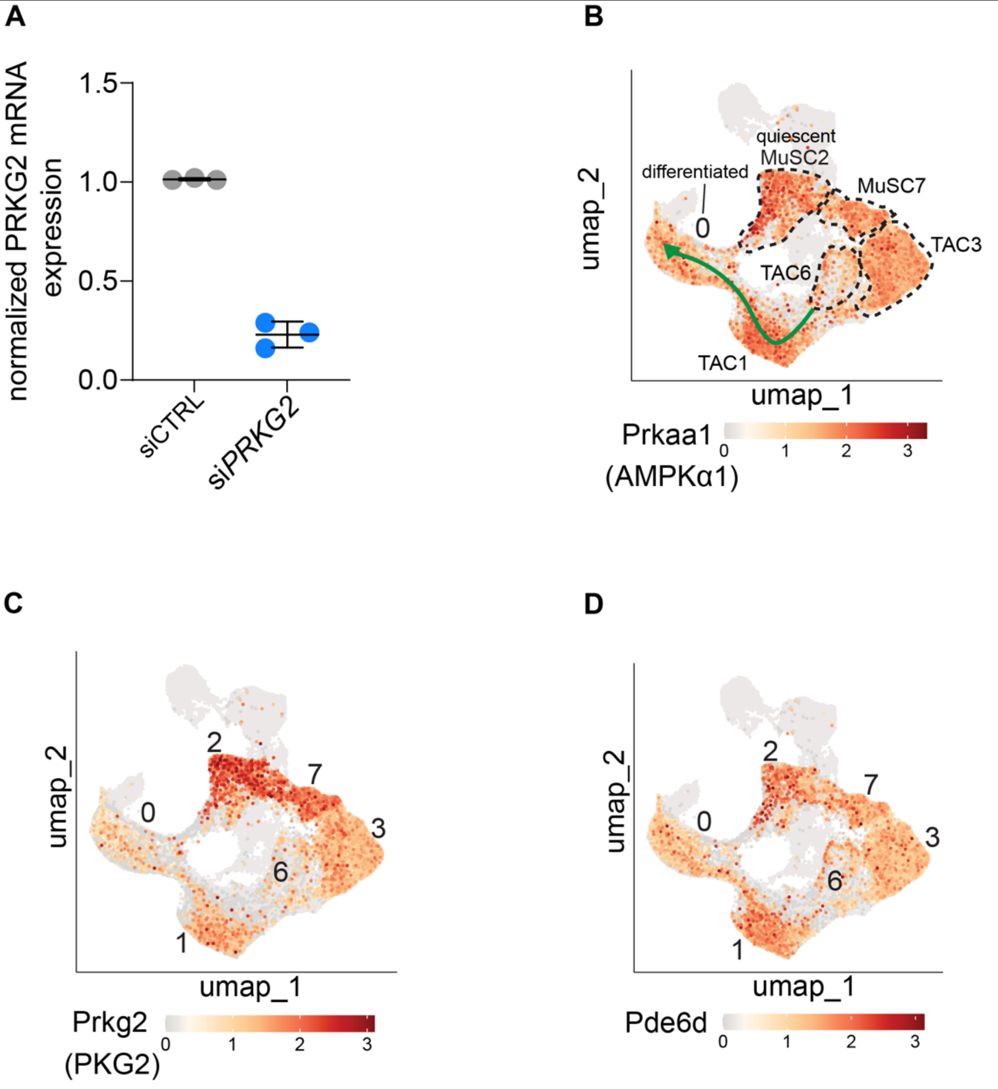


**Figure S3 related to Figure 6.**

(**A**) RT-PCR based *PRKG2* mRNA level quantification normalized to *GAPDH* mRNA of C2C12 cells grown under low serum for 72 h after transfection with siCTRL or si*PRKG2* (each 100 nM), N=3. Means ± SD are plotted. Statistical analysis was done with the unpaired t-test with Welch's correction.

(**B-D**) C2C12 cell differentiation scRNAseq UMAP representations of normalized gene expressions of *Prkaa1* (AMPKα1) (B), *Prkg2* (PKG2), (C) and *Pde6d* (D) derived from high and low serum samples. Relevant cell states and the differentiation trajectory are indicated in (B).
